# Evolutionarily Conserved Heat-Induced Chromatin Dynamics Drive Heat Stress Responses in Plants

**DOI:** 10.1101/2025.03.05.641738

**Authors:** Ming-Jen Yen, Kuan-Hung Lin, Lavakau Thalimaraw, Hsiu-Ru Yang, Kulaporn Boonyaves, Chia-Yi Cheng, Ting-Ying Wu

## Abstract

Eukaryotic organisms remodel chromatin landscapes to regulate gene expression in response to environmental stress. In plants, heat stress (HS) induces widespread chromatin changes, yet the role of heat-responsive Heat Shock Transcription Factors (HSFs) in chromatin remodeling and their evolutionary conservation remains unclear. Using chromatin accessibility profiling and transcriptomics in *Marchantia polymorpha hsf* mutants, we identify HSFA1 as a key determinant in positioning *cis*-regulatory elements (CREs) for HS-induced gene activation, a mechanism conserved across land plants, mice, and human cells. By integrating gene regulatory network modeling, we identify parallel transcription factor subnetworks, with MpWRKY10 and MpABI5B acting as indirect regulators of HS responses via phenylpropanoid pathways and general stress signaling. We further explore crosstalk between HS and abscisic acid (ABA) signaling, showing that while ABA modulates gene expression in an HSFA1-dependent manner, it does not induce broad chromatin remodeling, positioning it as a downstream regulator rather than a primary determinant of chromatin dynamics. To extend these insights, we develop a cross-species and cross-condition machine learning framework that accurately predicts chromatin accessibility and gene expression, demonstrating a conserved regulatory logic of stress-responsive chromatin and transcription dynamics. Our findings provide a conceptual framework for understanding how TFs coordinate chromatin architecture to drive stress adaptation in plants and potentially other eukaryotes.

## Introduction

Eukaryotes have evolved mechanisms to bridge genome activity and chromatin dynamics to environmental conditions, enabling adaptation to unfavorable growth environments. Environmental signals affect chromatin dynamics by regulating the binding of transcription factors (TFs) to specific DNA sequences known as *cis*-regulatory elements (CREs)^1^. Therefore, chromatin remodeling and dynamic changes are considered critical steps in gene regulation, including those occurring under heat-stress conditions (HS). The dramatic rewiring of gene and enhancer networks is driven by key TFs during dynamic environmental changes, such as heat shock transcription factors (HSFs) under HS conditions. These factors, in coordination with chromatin-modifying enzymes, remodel the chromatin architecture, thereby determining whether genes are activated or repressed in eukaryotic cells^2^.

HSFs underwent extensive duplications and functional diversification during the evolution of land plants^3^. They are classified into three subfamilies-HSFA, HSFB, and HSFC-based on their domain structures^4^. While HSFA1 is recognized as a conserved master transcriptional regulator of heat stress responses (HSR) in plants, HSFBs are thought to function as repressors or co-activators that not only regulate the HSR but also contribute to the regulation of developmental processes in plants^5,6^. In addition to transcriptional regulations, a recent study in tomatoes has shown that HS induces chromatin accessibility, with HSFA1a playing a crucial role in the dynamic formation of promoter-enhancer contacts in response to heat at several loci^7^. However, given the complexity of HSFs in plants, the general regulatory framework governing heat-induced chromatin accessibility and the extent to which HSFA and other HSF subfamilies contribute to this process remains unclear.

Open chromatin regions (OCRs) provide information about CREs, which encode the genomic blueprints for proper transcriptional patterns. CREs are often clustered together, forming *cis*-regulatory modules^8^. These modules facilitate dynamic interactions between TFs and their target binding sites, enabling the activation or suppression of gene expression. Consequently, the positioning and location of these regions underpin transcriptional variation spatiotemporally in response to environment, adaptation, and evolution^8^. It is well-established that HSFs bind to heat shock elements (HSEs) in the promoter regions of HSR genes^4^. However, recent studies showed that HSF1 binds not only to promoters but also to some intergenic loci in *Drosophila* and human cells during HSR^9,10^. Additionally, HSF1 binding near promoters does not always activate nearby genes^10^. These findings challenge the established relationship between TF binding at target sites and its role in transcription. Similarly, in *Arabidopsis*, HSFA1a binds to promoters and gene bodies of a few genes at high temperatures^11^. It is crucial to understand how temperature activates HSFA1 and other HSFs, leading to large-scale and rapid genome and transcriptome reprogramming. However, the mechanisms driving chromatin changes at temperature-responsive loci and their conservation in land plants remain largely unexplode.

HSRs form complex regulatory networks involving interaction between various TFs and stress hormones such as abscisic acid (ABA). Understanding whether such regulations operate in HSF-dependent or HSF-independent manners to help plants cope with high temperatures remains a long-standing question. For example, the circadian clock proteins REVEILLE 4/8 (RVE4/8) are essential TFs that function in parallel with HSFA1s to regulate the first wave of HS-induced gene expression and prepare plants for exposure to high temperatures during the day^12^. HSF-dependent regulations, on the other hand, may coordinate with subsequent chromosome structure changes and require co-regulation with other TFs to sustain a longer period of HSR^11^. While ABA plays a key role in abiotic stress tolerance, including drought, salt, and osmotic stresses, its role in HS is not straightforward^13^ as ABA signaling-deficient and ABA hyper-responsive mutants exhibit defects in both basal and acquired thermotolerance^14,15^. Additionally, ABA application was shown to repress heat-induced dephosphorylation of BRI1 EMS-SUPPRESSOR 1 (BES1), suggesting an inhibitory role for ABA in certain stages of the HSR^16^. This also highlights its role in expanding the HSR network in coordination with HSFA1a in Arabidopsis. Given the complete emergence of ABA signaling components and the diversification of HSFs in land plants, it is intriguing to investigate the extent to which HS and ABA signaling are coordinated, as well as how HSFs regulate ABA signaling at both chromatin and gene expression levels in an evolutionary context.

In this study, we profile the dynamics of chromatin accessibility and gene expression in each Mp*hsf* mutant under HS conditions in Marchantia. We found that MpHSFA1 is essential for HS-induced chromatin remodeling, while MpHSFB1 preferentially regulates chromatin accessibility under control conditions. We further demonstrated that HSFA1 expands its binding sites downstream of the TSS, a regulatory mechanism conserved across eukaryotes. Our multi-layered gene regulatory network reveals both HSFA1-direct and HSFA1-indirect regulatory pathways, with MpWRKY10 and MpABI5B acting as negative regulators that parallelly modulate the HSR in Marchantia. Additionally, we found that ABA affects gene expression independently of chromatin dynamics, suggesting its role is more downstream in stress signaling regulation. Using machine learning (ML), we accurately predicted gene expression across stress and species contexts, revealing that HS *cis*-regulatory codes are more diverse than ABA response codes. Together, our findings not only identify the role of HSFA1 in positioning CREs, which predict gene expression under HS conditions but also expand our understanding of how HSFA1-multilayered regulation facilitates adaptation to high temperatures.

## Results

### HS induces genome-wide chromatin remodeling in Marchantia

Stress-induced chromatin remodeling often leads to the rewiring of TFs regulation and ultimately affects morphological and phenotypical changes^17^. The observation of morphological defects under control conditions and the hypersensitivity to high temperatures in HSFA and HSFB double knockout mutants^3^ (refer to Mp*dko* mutants hereafter) (Figure 1A-1D) led us to hypothesize that HSFs are likely involved in chromatin remodeling under both normal and HS conditions in Marchantia. To capture the dynamics of HS-induced chromatin remodeling, we collected ATAC-seq and RNA-seq datasets from Tak-1 plants, Mp*hsfa1*, Mp*hsfb1*, and Mp*dko* mutants during HS treatment in Marchantia (Figure 1E). We identified ATAC-seq peaks across all Mp*hsfs* mutant libraries and observed that the chromatin dynamics changed globally after HS treatment (Figure 1F, Supporting Figure S1, Supporting Dataset S1). Genomic regions corresponding to these peaks are OCRs, which preferentially align with upstream (−1.5k bp from transcriptional start sites (TSSs)) regulatory regions (∼40% of the peaks, Supporting Figure S2A&S2B).

**Figure 1:**
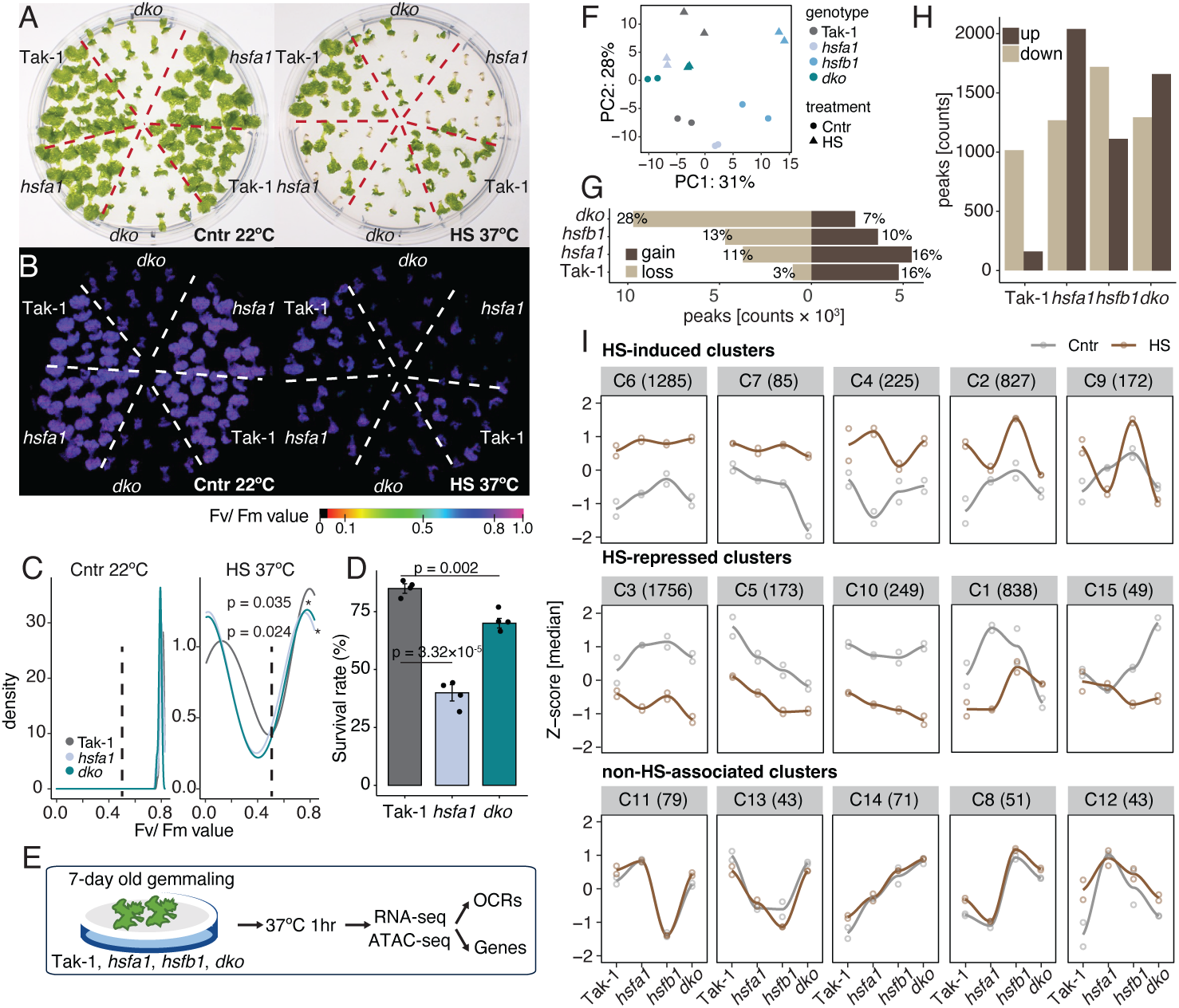
Chromatin accessibility undergoes dynamic changes under HS in Marchantia. A & B. Thermotolerance in 7-day-old Tak-1 plants, Mp*hsfa1*, and Mp*dko* mutants after a 90-minute HS treatment at 37°C in a water bath. Representative images of plants after five days of recovery are shown in GRB photos (A) and Fv/Fm photos (B). C. Density plots showing the distribution of Fv/Fm ratios under control conditions (22°C) and HS conditions (37°C). The vertical dashed line indicates the threshold (0.5) representing stress phenotypes. D. Bar graph displaying the survival rate (%) for each genotype after HS treatment. The p-value was determined by a two-tailed Student’s t-test; *n* = 60 gemmalings of each genotype across three independent experiments. E. Schematic of ATAC-seq and RNA-seq data collection for each genotype following HS treatment. F. Principal Component Analysis (PCA) comparing ATAC-seq biological replicates from each genotype under control and HS conditions. G. Bar graph showing the number of HS-induced gain or loss of peaks in each genotype. H. Bar graph displaying the number of differentially expressed (DE) open chromatin regions (OCR) in each genotype under HS conditions. I. Expression dynamics in hierarchical clusters of DE OCRs. Lines represent the average expression for each cluster, with the number of peaks in each cluster indicated in parentheses.

Differential analysis revealed 3% to 28% of regions that either gain or loss accessibility in response to HS in each genotype (Figure 1G). Among them, the Mp*dko* mutants showed the most lost peaks (28%), while the Tak-1 plants and Mp*hsfa1* mutants acquired the most gained peaks (16%) at HS conditions (Figure 1G & 1H), suggesting that mutant plants respond differently to HS-induced chromatin changes. Additionally, we examined the ATAC-seq signal in benchmarker genes such as Mp*sHSPs* and Mp*HSP90* at HS and found high chromatin accessibility in their promoter regions or gene bodies, while the signals were reduced in Mp*hsfa1* and Mp*dko* mutants (Supporting Figure S1). The distances of these peaks were enriched near TSSs (Supporting Figure S2A-S2C).

To gain the global view, we clustered the differential OCRs from at least one genotype under HS conditions into 15 groups based on OCR patterns (Supporting Dataset S2). Two-thirds of the clusters (10/15) were HS-associated, with clusters C2 and C9 showing decreased OCRs in Mp*hsfa1* and Mp*dko* mutants, suggesting that the opening of these regions is HSFA1-dependent. In contrast, the OCR patterns in one-third of clusters (5/15) were primarily genotype-dependent and non-HS associated, as the OCRs were not distinguishable between normal and HS conditions (Figure 1I). These findings suggest that HSFs are required for dynamic HS-induced chromatin changes.

### MpHSFA1 is required for HS-responsive genes with chromatin changes

We further examined whether changes in OCRs within each cluster corresponded to gene expression changes post-HS. We applied an orthogonal clustering approach to the RNA-seq data (referred to as rClusters hereafter) to establish correlations with our ATAC-seq clusters (Figure 2A, Supporting Dataset S3). Among the eight rClusters, 50% (4/8) contained heat-induced genes, which significantly overlapped with and were positively correlated with heat-induced aClusters, particularly C2 and C9 (FDR < 0.05; PCC > 0.5) (Figure 2B). These genes were downregulated in Mp*hsfa1* mutants and were functionally associated with HSR and protein folding (Figure 2C, left panel, Supporting Figure S2D). On the other hand, genes that were negatively correlated between HS-induced aClusters and rClusters (PCC < −0.5) were functionally associated with cell wall assembly (Figure 2C, left panel). In contrast, positively correlated repressed genes were linked to photosynthesis, while negatively correlated repressed genes were associated with lipid transport (Figure 2B & 2C, right panel, Supporting Figure S2D). Additionally, we found that several TF families were regulated at both the chromatin and RNA levels, as indicated by their intersection in Figure 2D (p < 0.05). These included MYB, AP2, WRKY, bHLH, and SBP. These findings suggest that chromatin dynamics play a crucial role in regulating HSR and TF gene expression in an HSFA1-dependent manner.

**Figure 2:**
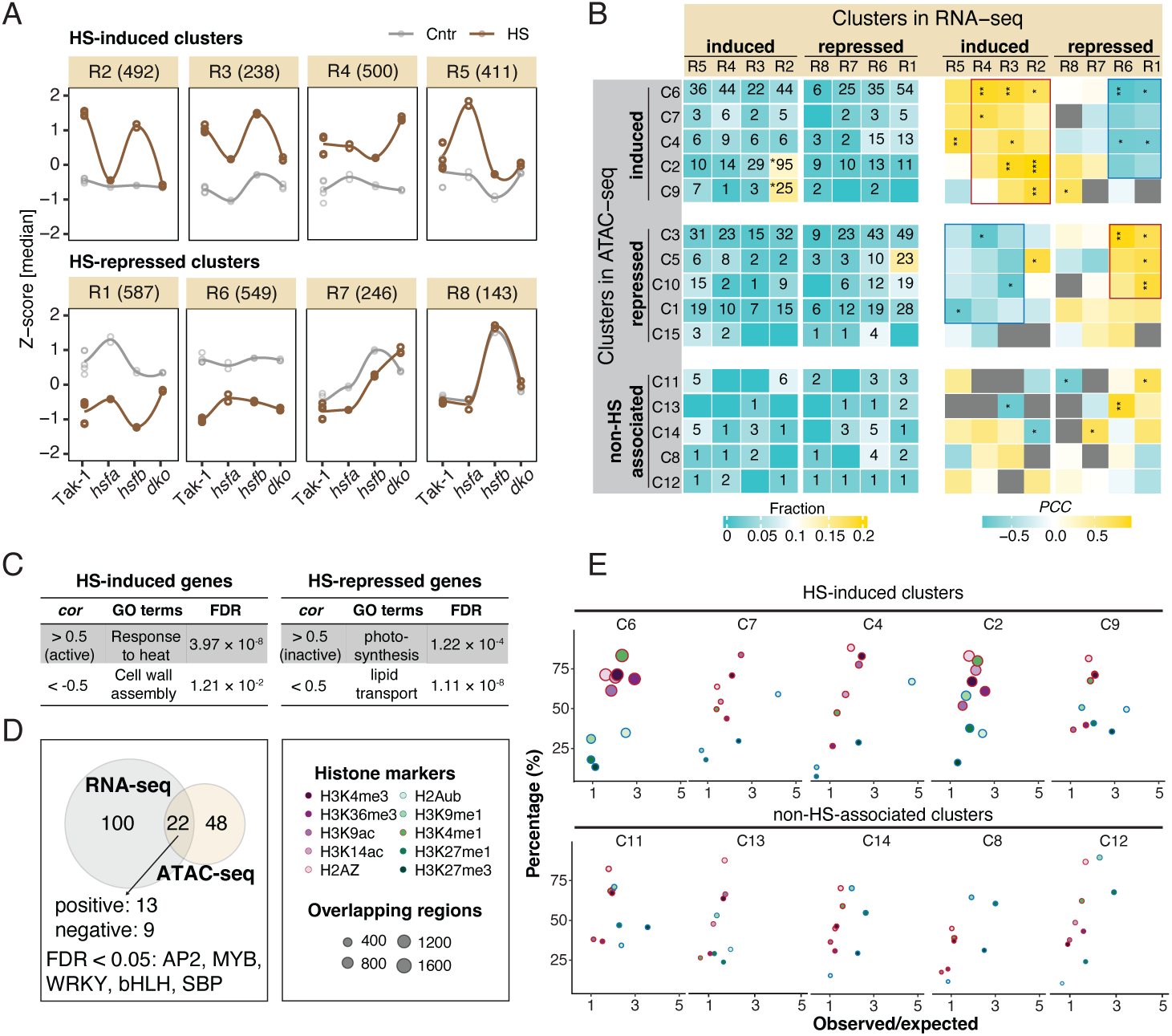
HSFA1 is required for HS-associated chromatin accessibility and gene expression in Marchantia. A. Expression dynamics in hierarchical clusters of differentially expressed genes (DEGs) across genotypes (rClusters). Lines represent the average expression for each cluster, with the number of genes in each cluster shown in parentheses. B. Heatmap showing the overlap of genes between rClusters and aClusters. Colors indicate the fraction of genes in rClusters found within aClusters. *p < 0.05, determined by Fisher’s exact test. Heatmap showing the similarity of expression patterns between rClusters and aClusters. Colors represent the Pearson Correlation Coefficient (PCC) between rClusters and aClusters, with significance levels denoted as *p < 0.05, **p < 0.01, and ***p < 0.001. C. Table of GO terms for genes in positively and negatively correlated groups from (B), respectively. D. Venn diagram showing TFs that are differentially expressed in both RNA-seq and ATAC-seq datasets (intersection). Significantly enriched TF families (FDR < 0.05) are listed, with FDR determined by Fisher’s exact test. E. The scatter plots illustrate the enrichment of different types of histone modifications in each cluster. Observed/Expected Ratio: This is calculated by dividing the observed overlap percentage by the background overlap percentage. The significance is assessed using the odds ratio. The observed overlap percentage indicates the proportion of differential OCRs that overlap with a specific histone modification marker within a given cluster, relative to the total number of differential OCRs in that cluster. The background overlap percentage is computed in a similar manner but is based on non-differential OCRs. The colors of the borders represent active (red) or repressive (blue) chromatin markers.

Epigenetic modifications on the chromatin are associated with gene expressions at HS^7^. To explore whether different clusters, particularly those activated under HS conditions, were enriched with specific histone modifications, we analyzed histone modification profiles using publicly available datasets from Marchantia^18^. We observed a significant enrichment (FDR < 0.01) of combinations of active epigenetic markers rather than individual markers in the clusters responsive to HS (Figure 2E, Supporting Figure S2E). These modifications were predominantly associated with the chromatin states related to promoter regions and gene bodies^18,19^. In contrast, both active and repressive epigenetic markers were more evenly distributed, with less enrichment in the clusters that were not associated with HS conditions (Figure 2E, Supporting Figure S2E). These findings suggest that HS-induced chromatin remodeling involves dynamic changes in multiple epigenetic modifications under HS conditions.

### HSFA1 is required for the positioning and reconfiguring of HS-related CREs

We used Transcription Factor Occupancy Prediction by Investigation of ATAC-seq Signal (TOBIAS)^20^ to analyze TF binding footprints in OCR clusters and to identify enriched CREs under HS in Marchantia (Supporting Figures S3&S4, Supporting Dataset S4). Enrichment was calculated for each cluster using non-regulated chromatin regions as controls (N), compared with proximal or distal OCRs (P). Both OCR types exhibited similar trends, with C2 and C4 showing the highest enrichment in CREs (Supporting Figures S3&S4). Co-occurring CRE analysis revealed dynamic pairing changes, with AP2EREBP-like CREs frequently co-occurring across all genotypes, while SBP-like CREs were specific to Mp*hsf* mutants (Supporting Figures S3C & S4C, Supporting Dataset S5). Combined with TF enrichment analysis (Figure 2D), these findings suggest that TF-CRE interactions coordinate distinct regulatory responses under HS, independent of HSFs.

Since MpHSFA1 regulates chromatin dynamics and gene expression under HS, we investigated its binding sites and locations. We found that enriched regulatory elements were primarily located downstream of the TSS in Tak-1 plants and Mp*hsfb1* mutants, whereas this distance was significantly reduced in Mp*hsfa1* and Mp*dko* mutants. This suggests that MpHSFA1 is essential for proper CRE positioning under HS in Marchantia (Figure 3A). To validate this, we first identified putative MpHSFA1 binding sites and target genes in Marchantia using DAP-seq (Supporting Figure S5A, Supporting Dataset S6). The TTCnnGAA binding motif was highly enriched in the MpHSFA1 dataset, closely matching known heat shock elements (HSEs) from HSFA1s in Arabidopsis and tomato (PCC > 0.9, e-value = 1.2×10^-350^) (Supporting Figure S5B & S5C). Genes with MpHSFA1 binding sites were primarily involved in canonical HSR or protein folding, similar to those in other species (Supporting Figure S5D & S5E). Next, we compared DAP-seq data from Arabidopsis^21^, tomato^7^, and our Marchantia dataset (Figure 3B, Supporting Dataset S6). Our analysis revealed that HSFA1 binding sites were enriched downstream of the TSS while binding sites for other HSFAs were concentrated near the TSS (Figure 3B). These results further support the notion that many HSFA1-dependent genes have transcriptional regulatory regions located downstream of the TSS. Additionally, we reanalyzed ATAC-seq data from tomatoes subjected to HS treatment for 1 and 6 hrs^7^. Enriched and highly ranked regulatory elements were predominantly located downstream of the TSS after HS treatment for 1 hr (Supporting Figures S5F and S5G).

**Figure 3:**
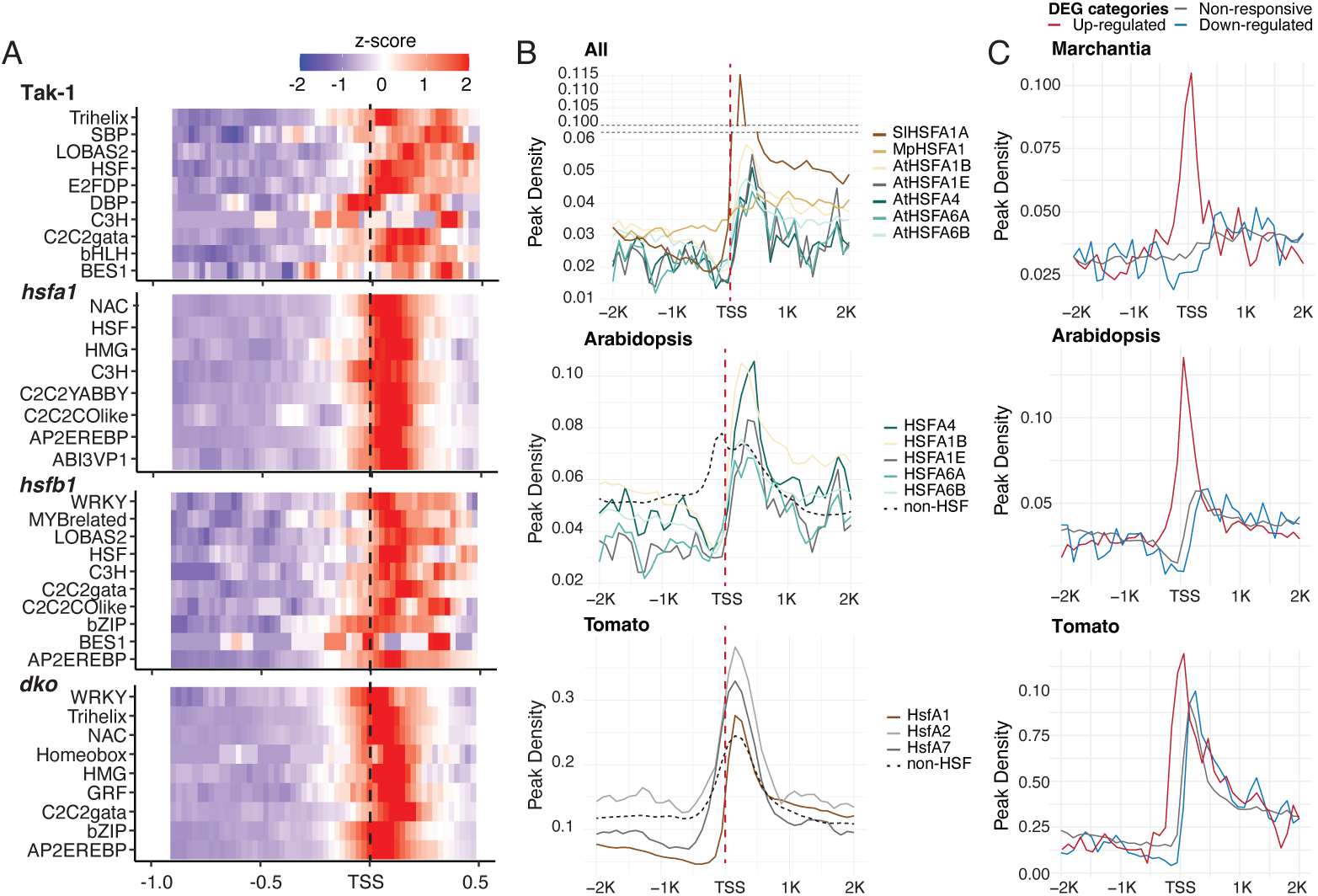
HSFA1 is required for the correct positioning of HS-related CREs. A. Distribution of top-ranked CREs (top 10 ranked TF binding motifs (TFBMs)) in each genotype. Site frequency represents z-scores for the ratio of CRE sites between positive and negative groups, calculated within a range from 1 kb upstream to 0.5 kb downstream of TSSs using a sliding window of 100 bp and a step size of 25 bp. B. Peak density plot (±2.0 kb) near TSSs for each DAP-seq dataset. Non-HSF refers to all other TFs in DAP-seq datasets, which served as background information. C. Peak density plot (±2.0 kb) near TSSs for MpHSFA1, AtHSFA1, and SlHSFA1 DAP-seq datasets of genes that were up-regulated, down-regulated, or non-responsive under HS. The DEGs in Arabidopsis and tomato were re-analyzed from previous publications^6,7^.

To explore the relationship between gene expression and HSFA1 binding sites, we divided the HSFA1-binding genes into three groups based on their expression patterns under HS conditions. The HSFA1 binding sites of HS-induced genes were primarily located near the TSSs, while those associated with down-regulated or non-responsive genes were preferentially found downstream of the TSSs (Figure 3C, Supporting Figure S6A&S6B). These patterns were not observed in *hsfa1* mutants, regardless of their gene expression patterns (Supporting Figure S6B). Further analysis on the function of orthologous genes in HSFA1-bound and HS-upregulated genes across four species showed that ∼13.5% of GO terms overlapped, primarily related to protein folding. In contrast, HSFA1-unbound and HS-upregulated genes exhibited more diverse functions, with no overlapping GO terms observed (Supporting Figure S6E&S6F, Supporting dataset S7). Consistent with this, binding sites of HSFA1 homologs in Arabidopsis (HSFA1a)^11^ and human (HSF1)^22^, as determined through *in vivo* ChIP-seq analysis, were also enriched in TSS downstream regions under HS conditions (Supporting Figure S6C). However, we observed a reconfiguration of HSFA1 binding sites toward the TSSs in HS-induced genes within the Arabidopsis ChIP-seq datasets. In contrast, this was not evident for HSF1 binding sites in the human datasets, as demonstrated by the distribution of binding patterns based on gene expression (Supporting Figure S6D). These findings suggest that while this regulatory mode is evolutionarily conserved in eukaryotes, the configuration of HSFA1 binding sites in HS-induced genes is more prominent in plants.

### MpWRKY10 and MpABI5B are required for the HSFA-indirect HS responses

Based on the analysis of distinct CRE enrichments and their combinations (Figure 3, Supporting Figure S3-S6) in different clusters and genotypes under HS conditions, we hypothesized that multiple regulatory mechanisms occur in the HSR in Marchantia, particularly those that bypass the regulation of HSFs. To further elucidate the relationship between these TFs-CREs in multi-layered HS regulation, we constructed a GRN that integrated various regulatory layers, including transcriptional regulation (RNA-seq), chromatin accessibility (ATAC-seq), and HSFA1-binding data (DAP-seq). To obtain a higher-confidence subnetwork, we selected nodes that contained the top 20% edges (top 20% based on what?) from the initial network, resulting in ∼1500 nodes, including 119 TFs. Of the 119 TFs incorporated into the GRN, 29 (∼25%) were bound by HSFA1 (Figure 4A, Supporting Dataset S7). When we applied the same GRN architecture to other Mp*hsfs* mutants, we observed significant changes in expression levels, with more TFs becoming less responsive to HS in Mp*hsfa1* and Mp*dko* mutants (Supporting Figure S7A). Furthermore, to understand the impact of HSFA1 binding and regulation, we divided all genes in the GRN into HSFA1-dependent and HSFA1-independent groups based on their expression differences between Mp*hsfa1* mutants and Tak-1 plants (Supporting Figure S7B). HSFA1-targeted genes showed a more significant change in expression within the HSFA1-dependent group (R = −0.27 vs. −0.18; −0.24 vs. 0.097) compared to non-HSFA1-targeted genes, while no significant change was observed in the HSFA1-independent group (R = 0.84 vs. 0.84) (Supporting Figure S7B).

**Figure 4:**
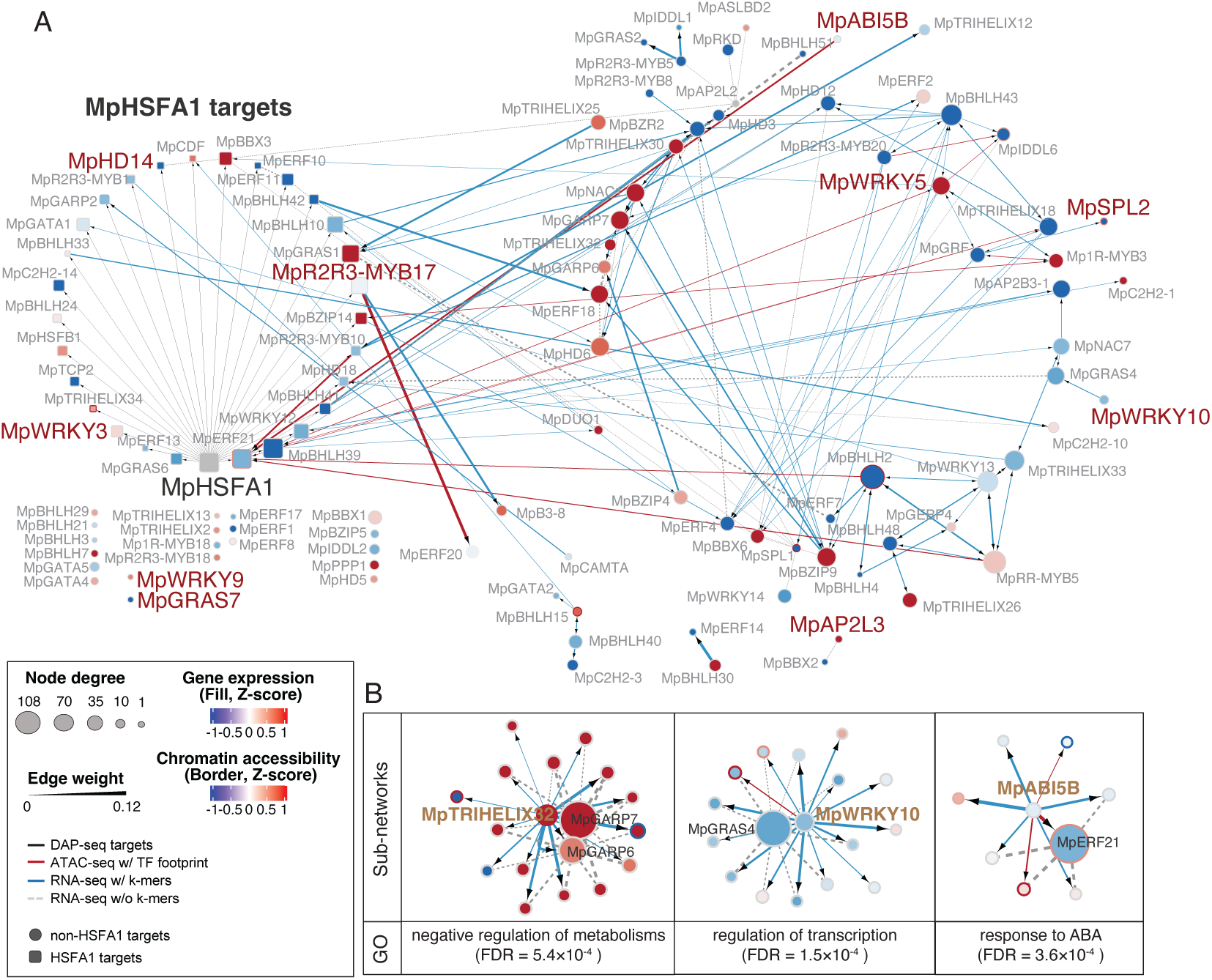
Multi-layered TF-GRN reveals HSFA1-direct and -indirect HS regulations. A. HSFA1-direct and -indirect co-expression network for HS-responsive genes in Marchantia. Square nodes represent HSFA1-binding target TFs, while circular nodes represent non-binding target TFs. Node colors indicate the relative change in expression (z-scores) of each gene across genotypes, and node size reflects the number of interacting partners in the network (see Methods). Edge width represents the significance of co-expression (see Methods). TF families discussed in the main text and/or selected for experimental validation are labeled in red. Black arrows indicate the direction of enriched CREs in each pair. B. Subnetworks of filtered nodes and their corresponding GO terms. These were selected based on their degree of centrality (bottom 25%), number of edges (greater than 25%), and proportion of CREs (greater than 50%) in the network.

To identify hidden regulators within the GRN, we specifically focused on several HSFA1-indirect (non-HSFA1-target) yet highly connected TFs for experimental validation (Figure 4B, Supporting Figure S7C). These TFs were selected based on their low degree of centrality (bottom 25%), a significant number of edges (greater than 25%), and a high proportion of CREs (greater than 50%) in the network. Subsequently, we focused on MpWRKY10 and MpABI5B because their subnetworks included GO terms related to transcriptional regulation and response to ABA (Figure 4B). Both Mp*wrky10* and Mp*abi5b* knockout mutants demonstrated greater tolerance to high temperatures compared to Tak-1 plants, as evidenced by higher survival rates (median: 32% vs. 60%, 28% vs. 60%, respectively) under HS (Figure 5A&5B, Supporting Figure S7D). Conversely, Mp*wrky3* knockout mutants exhibited a slightly lower but insignificant degree of thermotolerance compared to Tak-1 plants (Supporting Figure S7G). RNA-seq analysis revealed that *wrky10* mutants were less responsive to HS, as indicated by only about 57% of upregulated and ∼75% of downregulated DEGs compared to Tak-1 plants under HS conditions (Figure 5C&5D, Supporting Figure S7I). *wrky10*-specific HSR genes were significantly associated with phenylpropanoid pathways (q-value = 1.49 × 10⁻⁷) (Figure 5E). Conversely, *abi5b*-specific HS-responsive genes were more enriched in general stress response (q-value = 3 × 10⁻⁴) and hydrogen peroxide processes (q-value = 8 × 10⁻⁴) compared to those in Tak-1 plants (Figure 5F). Additionally, compared to *abi5b*- specific genes, *wrky10*-specific genes contained more differentially expressed TFs, including HSFA1-independent TFs within the HS-multilayer network (Supporting Dataset S8), despite having fewer HSR genes (Supporting Figure S7J). Our results suggested that these two TFs participated in HS regulation through HSFA-1 independent networks in Marchantia.

**Figure 5:**
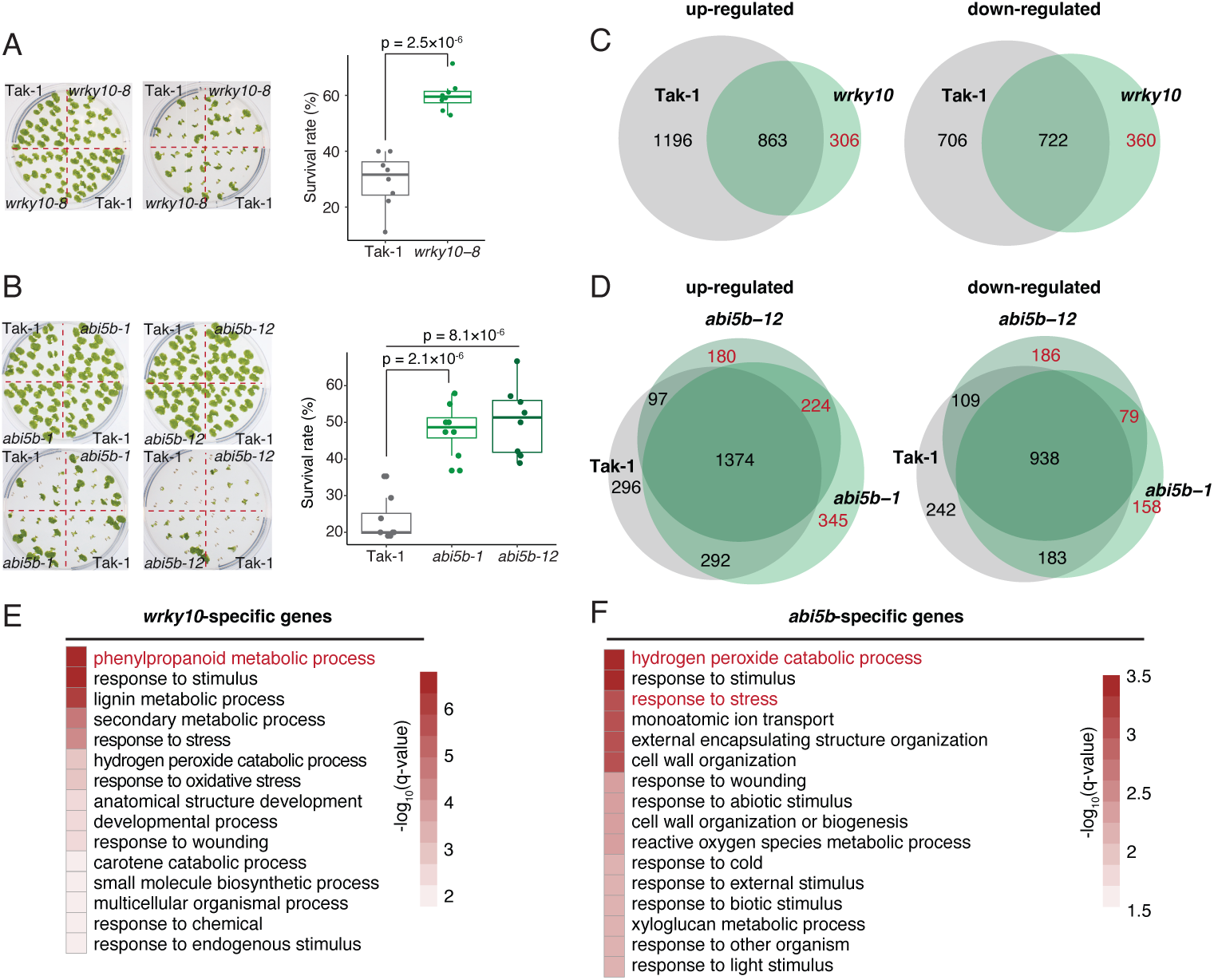
MpWRKY10 and MpABI5B are negative regulators of thermotolerance that function in parallel with HSFA1 in Marchantia. A. Thermotolerance in 7-day-old Tak-1 plants and Mp*wrky10* mutants after a 120-minute HS treatment at 37°C in a water bath. Representative images of plants after five days of recovery are shown in GRB photos. Box plots displaying the survival rate (%) for each genotype after HS treatment with the median values, the first and third quartiles and whiskers of maximum and minimum values. The p-value was determined by a two-tailed Student’s t-test; *n* = 20 gemmalings of each genotype across eight independent experiments. B. Same as those in (A.) but for the Mp*abi5b* and Tak-1 plants. The mutant information is listed in the Supporting Dataset S14. C. & D. Venn diagrams show the number of overlapping DEGs in *wrky10* vs. Tak-1 under HS (C.) and *abi5b* vs. Tak-1 under HS (D.). Genes differentially expressed in one *abi5b* line and with |log₂(FC)| ≥ 0.8 in the other were considered specific DEGs for further analysis. E. & F. Heatmaps display enriched GO terms of *wrky10*-specific (E.) and *abi5b*- specific (F.) upregulated genes.

### ABA does not induce intensive chromatin remodeling programs in Marchantia

Given that HS responses exhibit extensive crosstalk with hormone signaling pathways, such as ABA signaling^13^, we wondered whether MpHSFA1 is required for ABA regulation in Marchantia. To investigate this, we treated Tak-1 plants and Mp*hsfa1* mutants with varying concentrations of ABA for phenotypic observation (Figure 6A & 6B). The M*phsfa1* mutants displayed a significantly altered thallus area under ABA treatment compared to those grown under control conditions as early as 3 days post-treatment (Figure 6C, FDR < 0.01). However, the difference in thallus area between Mp*hsfa1* mutants and Tak-1 plants became insignificant after 5 days of treatment, suggesting that Mp*hsfa1* mutants are hypersensitive to ABA treatment. This led us to wonder if chromatin structure also responds to ABA in an HSFA1-dependent manner. To investigate this, we generated ATAC-seq and RNA-seq datasets for Tak-1 plants and Mp*hsfa1* mutants under ABA treatment (Figure 6D, Supporting Dataset S9). While ABA treatment triggered widespread transcriptional responses in both genotypes, surprisingly, we observed minimal changes in chromatin accessibility after ABA treatment, especially in Tak-1 plants (Figure 6D, Supporting Figure S8). This was further confirmed by the observation of 59 OCRs that gained accessibility and 54 OCRs that lost accessibility in response to ABA treatment in Mp*hsfa1* mutants, while no differential peaks were detected in Tak-1 plants (Supporting Figure S8). Given the minimal changes, we then focused on the analysis of 52 known ABA marker genes in Marchantia^23^. We examined whether chromatin accessibility correlated with gene expression changes under ABA treatment (Figure 6E, Supporting Dataset S10). These 52 marker genes showed a modest positive correlation between the ATAC-seq and RNA-seq datasets in both genotypes (*rho* = 0.28 and 0.30, *p* = 0.044 and 0.028 for Mp*hsfa1* and Tak-1 plants, respectively) (Figure 6E). However, a significantly positive correlation between chromatin accessibility and gene expression was observed only in Mp*hsfa1* mutants when comparing HS and ABA treatment but not in Tak-1 plants (*rho* = 0.31 vs. 0.06; 0.35 vs. 0.18 for ATAC-seq and RNA-seq, respectively, Figure 6F & 6G). These findings indicate that while ABA signaling triggers broad transcriptional responses, it does not induce substantial chromatin remodeling in Marchantia. Instead, chromatin accessibility changes appear to be predominantly governed by HSFA1, reinforcing its central role in orchestrating chromatin dynamics in response to environmental cues

**Figure 6:**
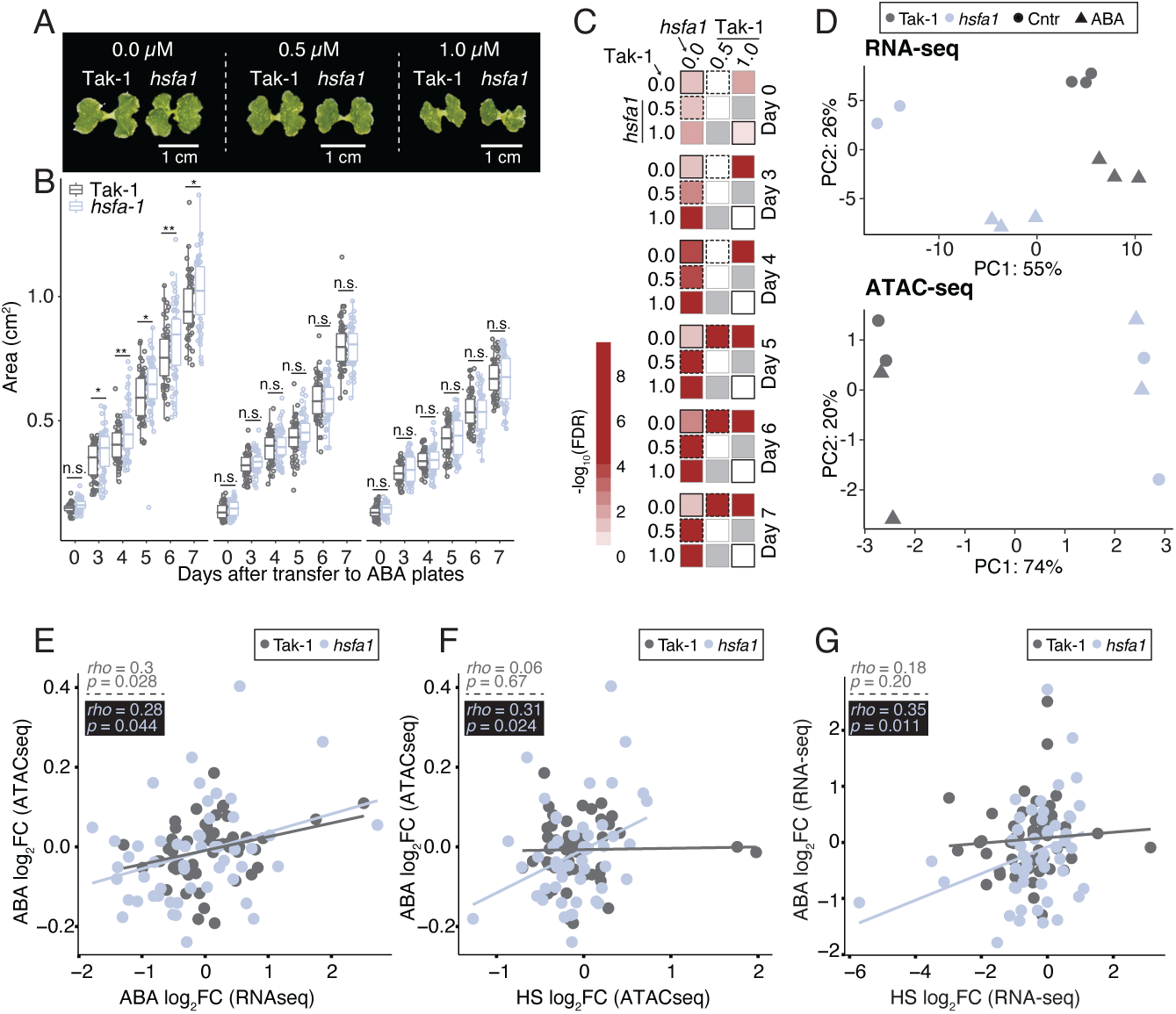
Mp*hsfa1* mutant is hypersensitive to ABA. A. Representative images of Mp*hsfa1* mutants and Tak-1 plants under control (mock, 20% EtOH) and ABA treatment conditions (0.5 or 1 µM ABA). Photos were taken seven days after ABA treatment on 1/2 × B5 agar plates. B. Thallus area of Mp*hsfa1* mutants and Tak-1 plants over time on 1/2 × B5 plates. Dots represent individual plant data, with *p < 0.05 and **p < 0.01, determined by the Mann–Whitney U test with Bonferroni correction. *n* = 30 plants per line. Boxplots display all data points (dots), median values, first and third quartiles, and whiskers indicating maximum and minimum values. C. Heatmap showing the significance of changes (−log10(FDR)) in each comparison over time, determined by two-way ANOVA. Grey cells indicate not applicable (n.a.) values. D. PCA comparing RNA-seq and ATAC-seq biological replicates from each genotype under control and ABA conditions. E. Scatterplots displaying the expression levels of 52 ABA benchmark genes between RNA-seq and ATAC-seq in Tak-1 plants and Mp*hsfa1* mutants. F & G. Same as (E), but comparisons are between HS and ABA ATAC-seq (F) and RNA-seq (G) datasets, respectively. The correlation was assessed using Spearman’s rank correlation coefficient.

### Species-specific predictability of HS and conserved ABA responses in land plants

We explored whether CREs information derived from OCRs and different sequence regions of genes could serve as predictive features in machine-learning models for HS-responsive gene expressions. First, we used enriched CREs (See Methods) identified from the 15 clusters, either using OCRs associated with genes or peaks alone for prediction. The AUROC scores across the 15 aClusters indicated that our model outperformed random guessing (0.5), with scores ranging from 0.67 to 0.92 for all clusters, except for clusters C8, C12, C14, and C15, where no enriched CREs were identified in the peak alone set (Figure 7A, Supporting Dataset S11). Next, we tested how CREs, identified using a combination of this information, could model the HS response. We identified CREs in the rClusters using promoter regions alone or combined with additional regulatory information from open chromatin sites (Figure 7B). The AUROC scores were higher in each group when additional regulatory information was included, ranging from 0.72 to 0.96 (Figure 7B, Supporting Dataset S11). Overall, these findings highlight that integrating ATAC-seq and RNA-seq data enhances the predictive power of HS gene expression models, underscoring the functional relevance of chromatin accessibility in transcriptional regulation under heat stress.

**Figure 7:**
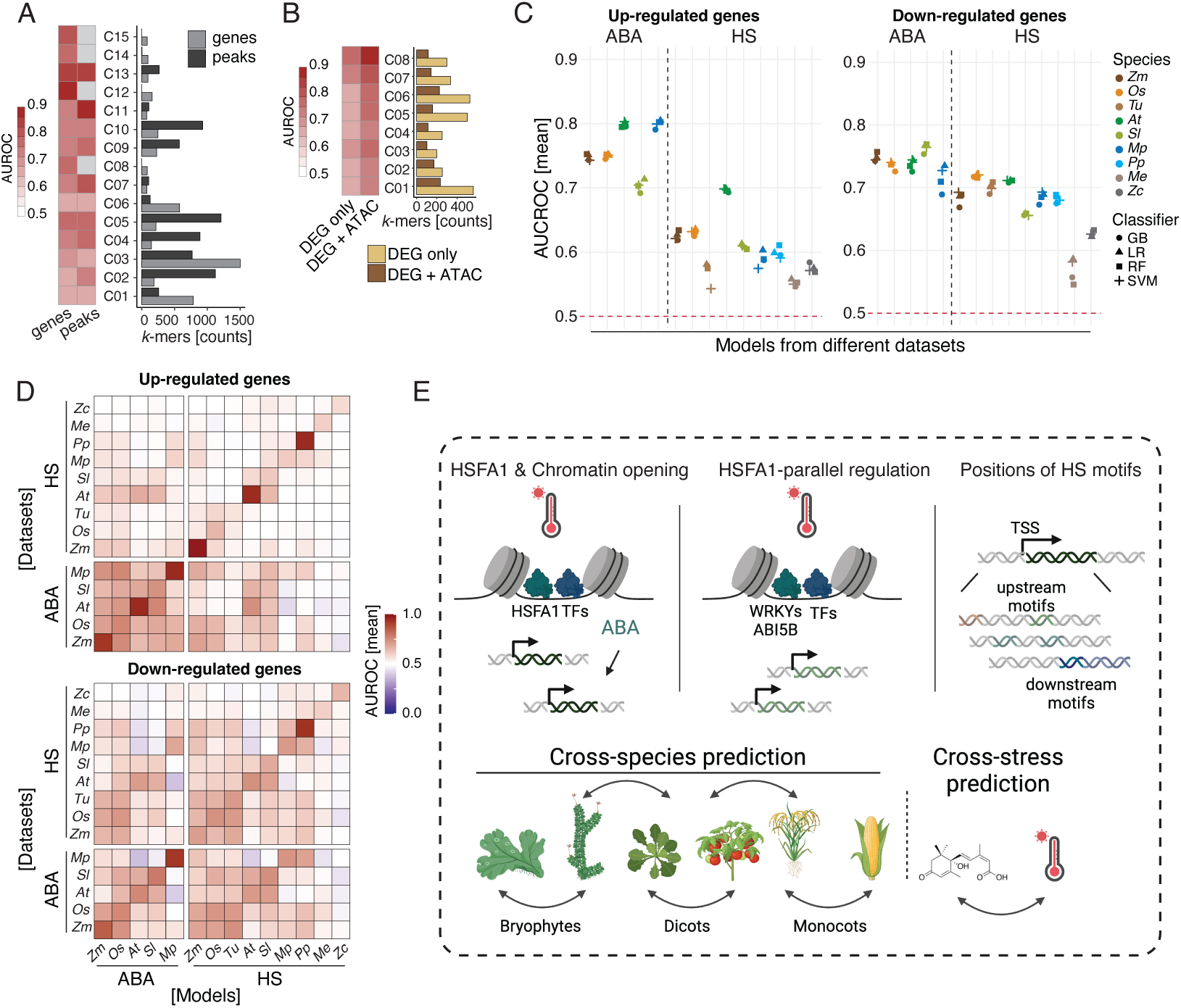
HS and ABA responses are predictable in Marchantia. A & B. Heatmaps showing ML performance for predicting HS responses using ATAC-seq datasets (A) or RNA-seq datasets (B) for each cluster in Marchantia. The AUROC scores represent the mean values from 10-fold cross-validation. Bar graphs indicate the number of enriched CREs identified in each cluster. C. Prediction performance, represented as AUROC (Area Under the Receiver Operating Characteristic) scores. Shapes indicate the mean AUROC scores for each ML algorithm, averaged over 10-fold cross-validation with standard deviation. The dashed line represents the baseline performance expected from random guessing. D. Heatmap showing cross-species and cross-stress prediction performance, using models trained on one species or one stress to predict HS or ABA responses in another. AUROC scores represent the mean values from 10-fold cross-validation. The full ML results and CRE information can be found in Supporting Dataset S10 & S12. E. The graphic summarizes the main findings of this study. We found that (I) HSFA1 is required for HS-induced chromatin remodeling, while ABA did not induce genome-wide chromatin changes. (II) ABI5B and WRKY10 are required for HSFA1-parallel HS regulation. (III) HSFA1 is required for the positioning of HS-related motifs in Marchantia. Additionally, we demonstrated that by using information from CREs, it is possible to predict gene expression across plant species and stress types.

To evaluate the broader applicability of our gene expression prediction framework across species and stress conditions, we reanalyzed publicly available RNA-seq datasets spanning algae, non-vascular plants, and vascular plants subjected to HS and ABA treatment^23–29^ (Supporting Dataset S12). The differentially expressed orthogroups (DEOG) and orthogroups of transcription factors (TFOG) showed significant similarities in the up-regulated groups, but not in the down-regulated groups under HS (Supporting Figure S9A & S9B). Most TF families were differentially expressed in more than one species within the TFOGs, indicating that transcriptional regulation is conserved during HS to a certain degree (Supporting Figure S9C). A similar trend was observed in the ABA response, where most DEOGs and TFOGs were expressed across multiple species (Supporting Figure S10). Based on this, we first constructed predictive models for each species under HS or ABA treatment (See Methods). The AUROC values ranged from 0.71 to 0.83 for ABA treatment predictions, while the AUROC values were lower for HS treatment predictions, ranging from 0.55 to 0.72 (Figure 7C, Supporting Dataset S12&S13). We then conducted cross-species and cross-stress predictions for HS and ABA treatments (See Methods). Predictions using ABA models generally outperformed those using HS models, regardless of whether predicting HS or ABA responses (overall mean AUROC ABA vs. HS = 060 vs. 0.57, Figure 7D, Supporting Dataset S12&S13). Interestingly, the AUROC values were significantly higher when using the HS model to predict ABA responses than when predicting HS responses (overall mean AUROC ABA vs. HS = 0.58 vs. 0.49 for up-regulated genes; 0.62 vs. 0.57 for down-regulated genes). Additionally, the AUROC scores were higher for land plants in both HS and ABA predictions (Figure 7D, Supporting Dataset S12&S13). These results suggest that HS-responsive CREs are more species-specific, while CREs involved in ABA responses may be partially conserved. Supporting this, we identified a greater number of highly ranked and frequently occurring CREs in the ABA dataset compared to the HS dataset across species (28 vs. 18), with 21% (n = 8) of CREs overlapping between the two stress conditions (Supporting Figure S11, Supporting Dataset S12&S13). Lastly, we evaluated the performance of cross-stress and cross-species predictions using the combinatorial information of OCRs and gene expressions (Supporting Figure S12, Supporting Dataset S12&S13). The prediction results for down-regulated genes showed substantial improvement in identifying HS-responsive genes across different species (overall mean AUROC = 0.62, Supporting Figure S12). Overall, our findings demonstrated the predictive power of CRE-based models in deciphering stress-responsive gene regulation, revealing a greater evolutionary conservation of ABA regulatory elements in land plants compared to the more species-specific nature of HS-responsive CREs.

## Discussion

Chromatin remodeling serves as the primary driver for initiating gene regulation in response to stress conditions or developmental processes^2^. It is also essential for various cellular functions, including the regulation of DNA replication, DNA damage repair, and cell cycle events^30^. Here, we uncover extensive HS-induced chromatin remodeling, revealing an orchestrated reprogramming of the Marchantia epigenome driven by HSFs. We identify regulatory regions of the genome that are specifically active or repressed in cells and uncover how the accessibility of these sequences is modified by signals that control thermotolerance in plants. We profiled genome-wide chromatin accessibility in each of the Mp*hsf* mutants exposed to HS (Figure 1). Our results led us to propose that HSFA1 is required for chromatin accessibility and the positioning of CREs under HS conditions, thereby regulating canonical HSR genes. On the other hand, HSFB1 maintains open chromatin of cell walls and development-related genes under control conditions (Figures 2-3). From the multi-layer GRN analysis, we identified several TFs, including Mp*WRKY10* and Mp*ABI5B*, that function alongside HSFA1 to regulate HS responses. Mp*WRKY10* regulates phenylpropanoid pathways, while Mp*ABI5B* is involved in general stress responses, respectively (Figures 4-5). In addition, Mp*HSFA1* regulates ABA responses by altering gene expression rather than modifying chromatin dynamics (Figure 6). Using the information from CREs, we accurately predict gene expression patterns under HS and ABA across species and found the *cis*-regulated codes in HS are more evolutionarily divergent (Figure 7). A graphic summary is provided in Figure 7E.

In the Mp*hsfa1* and Mp*dko* mutants, the expression of HSR genes correlated with the loss of chromatin accessibility (Figure 2). However, we still observed substantial chromatin opening events (∼16% and ∼7 % in Mp*hsfa1* and Mp*dko* mutants, respectively, Figures 1 & 2). Our analysis of the enriched CREs in these regions revealed an association with NAC, AP2/EREBP, ABI3/VP1, WRKY, and bZIP binding sites, suggesting that these TFs may function independently or in parallel with HSFA1 (Figure 3-5). Indeed, the GRNs, which integrate gene expression, chromatin accessibility, and HSFA1 binding data, reveal that a significant proportion of TFs (∼60% in the GRN) are not regulated by HSFA1 (Figure 4). While HSFA1 or HSF1 is generally considered the master regulator of the HSR, it has been shown that in mammalian cells, the majority of HSR genes are HSF1-independent with HSF1 binding to only a small subset of HS-induced genes^10^. Similarly, although HSFA1a is a primary factor that regulates the response to warm temperatures (27°C) in Arabidopsis, approximately 44% of HS-induced genes are not direct targets of HSFA1, and their regulation is not affected by HSFA1^11^. This indicates that heat-independent HSF1 binding, and possibly its function, is conserved in eukaryotes. Together with the analysis of co-expression with epigenetic markers under control conditions and co-occurring CREs (Supporting Figures S2-S4), our results suggest that HSFA1 may recruit chromatin-modifying proteins and other co-factors to target sequences, thereby increasing chromatin accessibility and affecting gene expression. Several such molecules, including SWI/SNF, H2A.Z, stress-inducible co-factors, and the RNA Pol II machinery, have been reported in Arabidopsis, human cells, and Drosophila^10,11,31–33^. Investigating the detailed reprogramming of these factors and determining whether they act in an HSFA1-dependent manner under different high-temperature conditions would be of great interest.

We observed HSFA1 binding sites distributed not only at TSSs but also within gene bodies in eukaryotes (Figure 3, Supporting Figure S5). This broad pattern of HSFA1 binding was not strictly associated with HS conditions to a certain extent (Figure 3, Supporting Figures S5&S6). Similarly, in human cells, HSF1 binding is largely decoupled from transcriptional activation. This decoupling is likely due to variability in transcriptional outcomes rather than differences in HSF1 binding and reflects the pervasive regulation of transcription by combinations of TFs^22^. Some low-affinity TF binding sites are known to be functional, with cooperativity among individual binding events driving genome-wide binding, including transcriptional enhancers^34,35^. Another possibility is that HSFA1 acts as a pioneer TF, where such factors do not restrict their binding locations. In this case, the chromatin targeted by HSF1 often resides in an open conformation even before HSF1 activation^27,36^. These two possibilities are not mutually exclusive, as they suggest that the local chromatin environment preconditions transcriptional responses and, together, enables HSFs and other TFs to orchestrate and access unique binding sites depending on the stress stimuli. In addition, the binding of HSF1 at remote genomic sites suggests potential functions beyond transcription (Figure 3, Supporting Figures S2-S5). A recent study utilizing 3D chromatin architecture analysis revealed that both proximal and distal CREs are enriched with HSFA1a binding motifs, suggesting that HSFA1a may promote or stabilize these interactions by maintaining contacts with promoter-distal regulatory element (enhancer) contacts under HS conditions in tomatoes^7^. This alternative regulatory mechanism may explain the slight shift, rather than a drastic reconfiguration, in HS-induced and HS-suppressed genes in tomatoes compared to other species (Figure 3, Supporting Figure S6). Nonetheless, the precise chromatin configuration that determines the accessibility of HSFA1 and other TFs to DNA, as well as the exact mechanistic controls underlying transcriptional outcomes at genes and distal or proximal CREs, remains to be explored.

ABA triggers changes in gene expression in Marchantia^23^, However, whether this transcriptional response relies on chromatin remodeling is still unclear. Our findings show that Mp*hsfa1* mutants display hypersensitive phenotypes and stronger gene expression, despite chromatin remodeling showing no significant changes in response to ABA treatment (Figure 6). In Arabidopsis, ABA induces extensive and dynamic chromatin remodeling in guard cells, roots, and mesophyll cells, with distinct patterns specific to cell types. Such processes are partially regulated by bZIP-type TFs^27^. The absence of clear ABA-induced chromatin changes in Marchantia could be due to cell-type-specific regulation masking the signals. Analysis of 52 ABA benchmark genes revealed a positive correlation between their expression levels under ABA and HS treatments (Figure 6). Consistent with this, CREs resembling bZIP binding motifs were among the top-ranked co-occurring CREs in Mp*hsfa1* mutants under HS conditions (Supporting Figure S3). This finding suggests potential cross-talk between these two signaling pathways^13^. The selected TFs for validation, WRKY10, and ABI5B, were also included among the 52 ABA marker genes (Figure 4, Supporting Dataset S10). However, the expression levels of WRKY10 and ABI5B did not vary between Tak-1 and Mp*hsfa1* plants under ABA and HS conditions. MpWRKY10 suppresses phenylpropanoid pathway enzymes and pigmentation genes in young Marchantia plants during phosphate deficiency^37^. The phenylpropanoid pathway was also highly up-regulated in Mp*wrky10* mutants under HS conditions (Figure 5), suggesting it serves as a common stress signaling cue for adaption to unfavored environments in plants^38^. On the other hand, MpABI5B is regulated by MpABI3, an evolutionarily conserved TF involved in ABA signaling in land plants^39^. Its role under HS and ABA treatments in our datasets (Figures 5 & 6) further highlights the conserved cross-talk between HS and ABA signaling.

CREs control gene expression, orchestrating stimulus responses and collectively influencing morphological changes^40^. Using CREs as a proxy to predict gene expression under ABA, HS stress, or cross-stress conditions in different species further supports the notion of evolutionarily conserved co-regulation between these two stresses (Figure 7). HS-induced genes were less predictable using CREs across multiple species, indicating that HS-associated CRE combinations evolved more rapidly, with species-specific CREs diversifying to help plants adapt to HS conditions (Figure 7). This is further supported by the identification of more GO functional groups during HS throughout land plant evolution^3^. Incorporating TF binding and chromatin accessibility data into our models improved their predictive performance (Figure 7). We observed that down-regulated genes were more accurately predicted across various stresses and multiple plant species (Figure 7, Supporting Figure S12). This aligns with previous studies indicating that most CREs responsible for repression are located in distinct sequence regions, primarily downstream of TSSs^41^. These results further provide the knowledge of regulatory grammar, which could represent the vocabulary of activating and repressing TFs and their combinatorial effects^40^. However, the models are not yet perfect, indicating that further investigation is needed to unravel the complexities of ABA and HS response perception, regulation, and the underlying molecular mechanisms^42^. Nonetheless, our findings highlight the intricate role of HSFs in regulating chromatin dynamics and the importance of CRE positioning. We also demonstrate that these regulatory mechanisms are conserved in eukaryotes under HS conditions.

## Material and method

### Plant growth conditions

*Marchantia polymorpha* wild type (accession Tak-1), Mp*hsfa1-13*, Mp*hsfb1-25* and Mp*hsfa1b1-56* (Mp*dko*)^3^, Mp*abi5b-1 &* Mp*abi5b-12,* Mp*wrky3-6*^43^ *&* Mp*wrky3-11*^43^ as well as Mp*wrky10-8*^37^ knockout mutants were grown on 1/2 × Gamborg B5 medium (pH 5.5) with 1% agar under continuous light (50–60 µmol m^-2^s^-1^) at 22°C.

### Generation knockout mutant lines

All cloning and transformation were performed as previously described^43^. Mp*abi5b* knockout mutants were generated using the clustered regularly interspaced short palindromic repeats/CRISPR Associated protein 9 (CRISPR/Cas9) method. Two gRNAs flanking the short DNA fragment of the target gene were designed and cloned into the pMpGE011 vector. Agrobacterium-mediated transformation of thalli was performed as previously described^44^, followed by the screening of transformants with 100 µg mL^-1^ cefotaxime and 0.5 µM chlorosulfuron. All experiments were conducted using the T2 generation. Detailed information about the Mp*abi5b* knockout mutants and gRNA sequences, as well as primers for sequencing, is provided in Supporting Figures S7 and Supporting Dataset S14.

### HS and ABA treatment in Marchantia

#### HS

Ten to twenty wild-type and mutant gemmae from the Tak-1 plants, Mp*hsfa1-13*, Mp*dko-56*, Mp*wrky3-6*, Mp*wrky3-11*, Mp*wrky10-8* and Mp*abi5b-1* & Mp*abi5b-12* lines were grown on 1/2 × B5 medium (pH 5.5) with 1% agar at 22°C under a 16-hour light/8- hour dark cycle for five days on cellophane. These 5-day-old plants were then transferred to treatment agar plates to grow for an additional two days. They were treated at 37°C for 90 or 120 minutes in a preheated water bath, followed by immediate transfer to a 22°C growth chamber with continuous light for recovery. Images were taken from Day 0 (immediately after treatment) to Day 10. The survival ratio was calculated as the green pixel area/total thallus area, where green pixels were defined by HSV values between [30, 20, 255] and [80, 255, 255]. Photographs and survival analyses were conducted on Day 5 post-HS. The Fv/Fm measurement was conducted as previously described^45^. Briefly, the plates were placed in the dark for 30 minutes before measurement with IMAGING-PAM (Walz, https://www.walz.com/).

#### ABA

Similarly, gemmae from Tak1 plants and Mp*hsfa1-13* lines grown on cellophane were transferred to treatment plates containing 1/2 × B5 medium (pH 5.5) with 0 (mock, 20% EtOH), 0.5, or 1 µM ABA. Plant images were taken every other day from Day 0 (immediately after treatment) to Day 7 to analyze the plant area. Morphological characteristics were extracted using OpenCV2 (https://opencv.org/).

### Collection of RNA-sequencing (RNA-seq) datasets and data analysis

#### Data collection

Marchantia gemmalings were grown in 1/2 × B5 liquid medium for seven days under continuous light and then subjected to HS at 37°C for 0 or 1 hour or treated with 0 (mock, 20% EtOH) or 50 µM ABA. Total RNA was extracted from the gemmalings using an RNeasy Plant Mini Kit (Qiagen, Hilden, Germany). Library preparation, including polyA enrichment and sequencing of 150-bp paired-end reads, was performed on the NovaSeq platform at GENOMICS, Taiwan.

#### Data analysis

Raw sequences from Marchantia were first processed to remove adaptors using fastp (v0.24.0). The cleaned reads were then mapped to the reference genome (Marchantia genome v6.1r1) using STAR (v2.7.9a) with default settings. Differentially expressed genes (DEGs) were identified using the R package DESeq2 (v1.42.1)^46^, with comparisons to expression levels at 0 hours (HS) or mock (ABA) conditions. A threshold of |log_2_(fold-change (FC))| ≥ 1 and a False Discovery Rate (FDR) ≤ 0.05 was applied using the “lfcShrink ashr” method. The DEGs were clustered based on their expression patterns, calculated from variance-stabilized transformation (VST)-transformed z-scores for the HS dataset (rClusters). Arabidopsis orthologs of Marchantia were identified as previously described^43^ for Gene Ontology (GO) analysis. GO term enrichment analysis was performed using the PlantRegMap GO enrichment tool (https://plantregmap.gao-lab.org/) with standard settings. Significantly enriched GO terms (q-value < 0.05, corrected for multiple testing using the Benjamini-Hochberg method) were summarized using REVIGO (http://revigo.irb.hr/) to remove redundant terms. The significantly enriched GO terms from each cluster were visualized as heatmaps using ComplexHeatmap^47^.

### Collection of Assay for Transposase-Accessible Chromatin with sequencing (ATAC-seq) datasets and data analysis

#### Data collection

The plants were grown and treated in the same manner as those used for RNA-seq data collection, and nuclei were isolated as previously described^18^. In brief, a total of 1.5 g of whole gemmalings were finely chopped with a clean razor blade in 4 mL of ice-cold Galbraith buffer (45 mM MgCl₂, 20 mM MOPS, 30 mM sodium citrate, 0.2% Triton X-100, and cOmplete Protease Inhibitor Cocktail) for 2-5 minutes in a chilled petri dish, followed by 10 minutes of incubation at 4°C. The lysate was then filtered twice through a 40 µm filter and transferred to an ice-cold 15 mL tube. Nuclei were pelleted by centrifugation at 500 × g for 10 minutes at 4°C, and the supernatant was carefully removed. The pellet was gently resuspended in 2 mL of Galbraith buffer and centrifuged again under the same condition. After removing the supernatant, the nuclei were resuspended in 1 mL of Galbraith buffer in a 5 mL round-bottom tube and stained with 10 µg/mL DAPI.

Flow cytometry was performed on a BD FACS Aria II using a 100 µm nozzle at a pressure below 20 psi. The photomultiplier settings were as follows: Forward Scatter (FSC) at 250 eV, Side Scatter (SSC) at 220 eV, and BV421 (DAPI channel) at 550 eV. Sorting gates were set based on FSC and SSC to exclude debris and potential doublets. Baseline fluorescence was established by running unstained control samples prior to sorting DAPI-stained nuclei. A total of 50,000 sorted nuclei were collected into 500 µL of Galbraith buffer (without Triton X-100) in 1.5 mL Eppendorf tubes. The nuclei were then pelleted by centrifugation at 1000 × g for 10 minutes at 4°C. The sorted nuclei were resuspended in a 50 µL tagmentation reaction mixture consisting of 25 µL of 2x TD buffer, 2.5 µL of TDE1 enzyme (Illumina Tagment DNA TDE1 Enzyme and Buffer Kits), and 22.5 µL of nuclease-free water. The reaction was incubated at 37°C for 30 minutes, followed by DNA purification with Zymo CHIP DNA Clean and Concentrator (Cat# D5205). The eluted DNA was amplified with dual-indexed Nextera primers using NEBNext 2x HiFi PCR Master Mix (New England Biolabs) for library construction.

For library amplification, 10 µL of transposed DNA fragments were mixed with 25 µL of NEBNext 2x HiFi PCR Master Mix, 6.25 µL of 10 µM T5 primer, and 2.5 µL of nuclease-free water. The corresponding T7 primer (6.25 µL) was added, and PCR amplification was performed with the following conditions: 72°C for 5 minutes (initial denaturation), 98°C for 30 seconds, then 5 cycles of 98°C for 10 seconds, 63°C for 30 seconds, and 72°C for 1 minute. Additional cycles were optimized via qPCR using PowerSYBR™ Green Master Mix on 5 µL of the amplified library. Amplified libraries underwent dual size selection using a 1.2x SPRIselect magnetic bead ratio (Beckman Coulter). Library quality was assessed with a Bioanalyzer High Sensitivity DNA Analysis kit (Agilent) at the Genomic Technology Core, IPMB, Academia Sinica, Taiwan. ATAC-seq libraries were sequenced in PE150 mode on a NovaSeq 6000 at GENOMICS, Taiwan.

#### Data analysis

Raw ATAC-seq sequence data were trimmed to remove adapters and were quality-checked using fastp (v0.24.0) and FastQC (v0.11.9). Processed reads were aligned to the Marchantia reference genome (v6.1r1) with Bowtie2 (v2.4.4). Alignments with low quality (mapping quality < 2), those mapped to mitochondrial or chloroplast genomes, and PCR duplicates were identified and removed using Samtools view (v1.14). ATAC-seq peaks were identified across biological replicates with MACS2^48^ (v2.2.7.1) using the following parameters: -f BAMPE -g 1.6e7 --cutoff-analysis --nomodel --shift - 100 --extsize 200. Identified peaks were consolidated using BEDtools merge to generate a consensus peak list. ATAC-seq alignments were assigned and quantified at these peak regions using the featureCounts function from subread (v2.0.6), and the resulting data was used for subsequent analyses. Differential chromatin accessibility analysis was conducted using DESeq2 (v1.42.1), with regions identified by a threshold of |Log_2_FC| ≥ 1 and an FDR < 0.05, applying the “lfcShrink ashr” method. Accessibility z-scores were computed from VST-transformed values, and hierarchical clustering was performed with the fastcluster package in R (aClusters). Genomic regions were defined and annotated using the R packages GenomicFeatures^49^ (v1.54.4), GenomicRanges^49^ (v1.54.1), and ChIPseeker^50^ (v1.38.0) with the default setting. For data visualization in the Integrated Genome Browser (IGV), bigwig coverage tracks were created from merged replicate BAM files using deepTools^51^ (v3.5.5) bamCoverage with a 10-bp bin size and Reads Per Kilobase Million (RPKM) normalization. Regions were linked to the closest TSS in the Marchantia genome for gene assignment. GO term enrichment analyses were conducted using the same protocol as for RNA-seq.

### Analysis of differential binding *cis*-regulatory elements (CREs)

TOBIAS (v0.13.3) was used to identify the binding positions of individual CREs within differential open chromatin regions (OCRs)^20^. The TOBIAS ATACorrect module was applied with the “--peaks” parameter set to all differential OCRs. Subsequently, the ScoreBigwig function was executed with the “--regions” parameter set to the differential OCRs. TOBIAS BINDetect module was then performed using the HS and control groups for each genotype according to its manual^20^. In brief, CREs were retained for further analysis if their binding events had a −log_10_(p-value) exceeding the 95^th^ quantile and/or differential binding scores above the 95^th^ quantile. Proximal and distal ATAC-seq CREs were analyzed separately. For each CRE, the median binding strength (log_2_FC of HS vs. control) and the total number of binding sites were calculated within each aCluster. The top 10 binding events were ranked based on the ratio of treatment-specific sites to total sites and further prioritized by their median binding strength.

### Analysis of CREs co-occurrence

TF-COMB (v1.1)^52^ was employed to analyze CRE co-occurrence. Briefly, the market basket algorithm was applied to identify co-occurrence rules (pairs of specific differential CREs) for each genotype. To determine significance, cosine similarity and z-scores within the top 95% quantile were calculated. The relationships among significant rules were visualized as a network using Cytoscape (v3.10.0), with cosine similarity representing the strength of co-occurrence as edges.

### DNA Affinity Purification Sequencing (DAP-seq) and data analysis

#### Protein Expression and Purification

The coding sequences (CDS) for Marchantia HSFA, and the N-terminal of HaloTag, along with the sequences for the SP6/T7 promoter and T7 terminator, were synthesized by Gene Universal, USA. These sequences were sub-cloned using BbsI into pAGM9121 as level 0 modules and further assembled via Golden Gate cloning into the level 1 expression vector pICH47732 to generate an SP6/T7 promoter-driven CDS with a N-terminal HaloTag and a T7 terminator. Recombinant proteins were expressed using the TNT SP6/T7 High-Yield Wheat Germ Protein Expression System (Promega) following the manufacturer’s protocol. Briefly, 2.5 μg of plasmid was used to translate protein *in vitro* at 30°C for 2 hours using a reagent mixture. Protein purity and integrity were confirmed by analyzing 10% of the final mixture via SDS-PAGE confirmed protein purity and integrity.

#### DNA Fragmentation and Preparation

Approximately 1 g of 14-day-old Marchantia gemmalings were collected and homogenized under liquid nitrogen. Genomic DNA was extracted using the phenol/chloroform method and sheared mechanically to an average size of 100-500 bp. The DNA fragments were end-polished, A-tailed, and ligated with a truncated Y-shaped adapter to prepare them for the protein-DNA binding assay.

#### Formation of Protein-DNA Complexes

*In vitro* translated proteins were incubated with HaloTag beads (Promega) for 1 hour at room temperature and purified using a magnetic separator with three washes in binding buffer (PBS with 0.1% NP-40). The purified beads-TF complexes were then incubated with the sheared, adapter-ligated DNA in the binding buffer to facilitate protein-DNA interactions. The mixture was incubated at room temperature for one hour to form TF-DNA complexes. Bead-TF-DNA complexes were captured using a magnetic base and washed three times to remove non-specifically bound DNA. Bound DNA was eluted from the beads by heating the mixtures to 98°C for 10 minutes. PCR was used to enrich the protein-bound DNA, incorporating a unique index for each sample to prepare them for multiplex sequencing. DNA fragmentation, adapter ligation, and library preparation were performed by the Genomic Technology Core at IPMB, Academia Sinica. The DAP-seq libraries were sequenced in PE150 mode on a NovaSeq 6000 at GENOMICS, Taiwan.

#### Data analysis

Raw DAP-seq reads were processed to remove adapters and perform quality control using fastp (v0.24.0) and FastQC (v0.11.9). The reads were then aligned to the reference genome (Marchantia genome v6.1r1) using Bowtie2 (v2.4.4)^53^ with the following parameters: --mp 6 -L 22 --dovetail. Low-quality alignments (mapping quality < 2), reads aligned to mitochondrial and chloroplast genomes, and PCR duplicates were flagged and removed using the Samtools view function (v1.14). DAP-seq peaks were called using the MACS2 callpeak function with the following parameters: -f BAM -g 1.6e7 -B -q 0.05 -m 2 50. Bigwig files were generated from merged DAP-seq BAM files using the deepTools (v3.5.5) bamCoverage function with a bin size of 10 bp, normalized via RPKM. To map peaks to target genes, DAP-seq peaks from each replicate were processed individually. The ‘ClosestGene’ method from the TFTargetCaller package (v0.7)^54^ was used to calculate target scores and q-values for all protein-coding genes annotated in the Marchantia reference genome (v6.1r1). Only those present in at least two replicates were retained. Genes found in all replicates were included in the GO term analysis.

### Construction of gene regulatory networks (GRNs)

Normalized gene expression counts were used to construct GRNs. *k*-mers (CREs) associated with each group were identified using the *k*-mer pipeline^55^, followed by calculating their Pearson correlation coefficients (PCC) with motifs in the Arabidopsis DAP-seq database^56^. The motif family with the highest PCC was selected to represent each *k*-mer. GRNs were generated using ARACNe-AP^57^ and GENIE3^58^, with TFs assigned before network construction. In the ARACNe-AP network, only edges with Bonferroni-corrected p-values below 0.05, based on mutual information (MI) values, were retained. For GENIE3, a baseline edge weight of 0.01 was set, and the interquartile range (IQR) of all edge weights was calculated. The third quartile (Q3) edge weight was used to filter high-confidence subnetworks. An intersection GRN was created by merging the outputs of both algorithms. To form a representative subnetwork, the top 20% of edges ranked by weight and MI value were selected. Nodes were grouped before constructing each GRN, integrating enriched *k*-mers and motif families. Edges were labeled as “with *k*-mer” if *k*-mers associated with hub TFs were found in the promoters of target nodes (indicating a “direct-linkage” relationship); otherwise, they were labeled as “without *k*-mer.” Similarly, edges were labeled “with TF footprints” if *k*-mers associated with hub TFs were present in the OCRs of target nodes. GRNs were visualized using Cytoscape (v3.10.0).

### Cross-species and cross-stresses prediction, and reanalysis of public datasets

#### RNA-seq re-analysis

Details on accession numbers and sources of publicly available datasets are summarized in Supporting Dataset S11. Raw FASTQ files were retrieved using SRA toolkits (v3.3.0). Raw reads were trimmed with Trimmomatic (v0.32) to remove low-quality bases and adapters. RNA-seq reads were then aligned to the respective reference genomes for each species using HISAT2 (v2.2.1), with detailed genome information and versions provided in Supporting Dataset S12. Read counts per gene were obtained using featureCounts from the Subread package (v2.0.7). Differential expression analysis was conducted with the DESeq2 package (v1.42.1). Genes were classified as DEGs if they had llog₂FC| ≥ 1 and FDR < 0.05, while genes with |log₂FC| < 0.2 were designated as negative control genes (TN).

#### DAP-seq and ChIP-seq re-analysis

Publicly available datasets were retrieved as mentioned and summarized in Supporting Dataset S11. After trimming, reads were mapped to the reference genome using Bowtie2 (v2.4.4). Peaks were identified using MACS2 (v2.2.7.1) for peak calling with default settings. From each peak summit, a 50 bp flanking DNA sequence was extracted and analyzed for motifs using MEME (v5.4.1)^59^, with a specified motif width of 6 to 20 base pairs. Peaks overlapping promoter regions (defined as −1000 to +500 bp relative to the TSS) were classified as DAP-seq or ChIP-seq peak-associated genes.

#### ATAC-seq re-analysis

To identify chromatin regions accessible under stress conditions, we analyzed publicly available data from ATAC-seq datasets, with detailed information in Supporting Dataset S11. After trimming, reads were mapped to the reference genome using Bowtie2. For each dataset, under both control and stress conditions, MACS2 was used to identify accessible chromatin regions (v2.2.7.1). All identified peaks were then merged across conditions using “bedtools merge”. Counts for each peak were obtained with “bedtools multicov” to enable quantitative comparison between conditions. Differentially accessible peaks were identified using DESeq2. Peaks with an FDR < 0.01 and a log₂FC > 1 were classified as stress-induced open peaks, while those with |log₂FC < 0.2| were designated as negative control peaks (TN).

#### Stress-associated CREs identification with machine learning (ML) prediction

To identify stress-associated CREs, we employed an ML approach using CREs as input features. Genes up- or down-regulated under stress conditions were treated as positive examples (TPs), while negative control genes served as TNs. For each gene, CRE counts were calculated within promoter regions (−1000 to +500 bp relative to the TSS), with CRE lengths ranging from 6 to 12 base pairs. To prevent duplication, the reverse complement of each CRE was considered, allowing each unique sequence to be represented only once. To evaluate the statistical significance of CRE distribution differences between TPs and TNs, we performed the Wilcoxon rank-sum test with Bonferroni correction. The 100 most significant CREs, determined by the smallest adjusted p-values, were retained for model training. Additionally, Cohen’s d was calculated for each significant CRE to assess effect size, quantifying the difference between TP and TN groups. For each significant CRE, we also calculated its positional distribution across promoter regions, and log₂FC of CRE counts between TP and TN genes was computed for each position, allowing further exploration of positional relevance. Our ML pipeline, adapted from a previous study^60^, was implemented using scikit-learn version 1.5.0 in Python 3.10.14. The dataset was divided into training (80%) and testing (20%) subsets to ensure robust model evaluation. To evaluate predictive efficacy, we constructed models using four different algorithms—Random Forest (RF), Gradient Boosting (GB), Logistic Regression (LR), and Support Vector Machine (SVM). Model optimization involved a grid search to identify the best hyperparameter settings for each algorithm, with performance validation through 10-fold cross-validation. The area under the curve for the receiver operating characteristic (AUROC) score served as the primary metric for selecting the best model for each dataset. Models with the highest AUROC values were further tested on a separate validation set, confirming their generalizability and robustness in predicting stress-associated genes.

#### CRE-TF assignment

CREs were first converted into MEME format to enable TF assignment. To identify the closest matching TFs, Tomtom from the MEME suite^59^ was used to compare CREs against the Arabidopsis DAP-seq database^21^, and TFs with the lowest e-values were assigned.

### Prediction of OCRs and gene expression

#### Feature collection

We identified CREs enriched in the regulatory (or peak) regions of differentially expressed gene/peak (DEG/DEP) clusters. For RNA-seq, regulatory regions were defined as 1 kb upstream to 0.5 kb downstream of a DEG’s TSS. For ATAC-seq, the peak region was determined based on peaks identified using MACS2. The process for identifying CREs followed a previously described method^55^. Briefly, the analysis began by evaluating all possible 5-mer oligomers for enrichment in DEG/DEP clusters, retaining those with significant enrichment (adjusted p-values < 0.05, determined using Fisher’s exact test with Benjamini-Hochberg correction for multiple testing). This procedure was repeated iteratively until no additional enriched extended *k*-mers were identified. When two significantly enriched *k*-mers overlapped, only the one with the lower adjusted p-value was retained.

#### ML prediction

We used the ML pipeline, as mentioned above. For each aCluster and rCluster, we split balanced data into training (80%) and testing (20%) sets and tested three classification methods: RF, SVM, and LR. We used ten-fold internal cross-validation to select the optimized hyperparameters. AUCROC score was used to select the best model for each cluster. The negative dataset was defined as non-differentially expressed OCRs with |log_2_FC < 0.8| (n = 10092) across all groups.

### Orthologous analysis

Whole proteome sequences of *Z. circumcarinatum*, *M. endlicherianum*, *A. thaliana*, *O. sativa*, *S. lycopersicum*, *Triticum urartu*, *Z. mays,* and *M. polymorpha* were downloaded in FASTA format from the following databases: *Z.circumcarinatum* and *M. endlicherianum*: Zygcir6981b_2 and Mesen1_1 from JGI PhycoCosm (https://phycocosm.jgi.doe.gov/phycocosm/home); *A. thaliana*: Araport11 from TAIR (https://www.arabidopsis.org/), *O. sativa*, *S. lycopersicum,* and *Z. mays*: MSUv7.0, ITGA5.0, Zm-B73-REFERENCE-NAM-5.0 from Phytozome (https://phytozome-next.jgi.doe.gov/), *T. urartu*: Tu2.1 (https://www.mbkbase.org/Tu/) and *M. polymorpha*: v.7.1 from MarpolBase (https://marchantia.info/). The proteome sequences from primary transcripts were used to construct OrthoFinder (v3.0)^61^ groups. The genes assigned to the OrthoFinder groups were used for further analysis. The detailed genome version and source are listed in Supporting Dataset S12.

### Accession numbers

The gene IDs for the mutants used in this study are as follows: HSFA1 (MpVg00470); HSFB1 (Mp4g12230); WRKY3 (Mp5g05560); WRKY10 (Mp7g06550); ABI5B (Mp2g22820).

## Data availability

Raw sequencing data have been deposited into the National Center for Biotechnology Information Gene Expression Omnibus under accession numbers (GSEXXXX).

## Code availability

The custom code used for this paper is available on GitHub (https://github.com/LavakauT/HS_ABA_script).

## Conflict of interest

The authors declare no competing interests.

## Author’s contribution

TYW conceptualized and oversaw all the experiments and analyses. LT performed preliminary analyses. MJY and TYW performed formal bioinformatic analyses with assistance from CYC. LT performed all sequencing experiments with the assistance of SRY, KHL, and TYW. KB and KHL designed the constructs for CRISPR/ Cas9 editing mutants. KHL and SRY performed DAP-seq, mutant generation, and physiological assay. TYW wrote the manuscript with assistance from all authors. KHL, KB, CYC, and TYW edited the manuscript.

## Acknowledgment

This study is supported by funding from the National Science and Technology Council (NSTC) 112-2311-B-001-009-MY3 and Academia Sinica (AS) AS-CDA-113-L03 provided to TYW. We thank Dr. Daisuke Urano (Temasek Life Sciences Laboratory and University of Singapore, Singapore) for kindly providing Mp*hsfa1*, Mp*hsfb1*, Mp*dko*, Mp*wrky3,* and Mp*wrky10* mutants. We thank Ms. Shu-Jen Chou and Ms. Mei-Jen Fang from the Genomics Core at IPMB for preparing ATAC-seq and DAP-seq sequencing libraries and performing Sanger sequencing, respectively. We also thank the Flow Cytometry Core at IPMB and IBMS, AS, for assisting with nuclei sorting. Additionally, we acknowledge the Bioinformatics Core at IPMB for providing high-performance computing support. Lastly, we thank Dr. Ming-Jung Liu (ABRC, AS) to critically read the manuscript.

## Supporting information

**Supporting Figure S1 HS-induced chromatin remodeling in Marchantia.** A. Heatmap of ATAC-seq signal (log_10_(RPKM+1)-normalized counts) centered on ATAC-seq peaks (±3.0 kb) showing HS-regulated chromatin accessibility across clusters and genotypes. B. Genome browser snapshots for three representative genes (MpGST26 - Mp2g026510, MpHSFP90-4 - Mp5g15940, and MpHSP17.8A1 - Mp7g07900) displaying changes in ATAC-seq signal in upstream regions and gene bodies following HS treatment. Scale bars represent 500 bp, and gray-shaded boxes indicate regions with differential accessibility.

**Supporting Figure S2 Characterization of OCRs based on the distance from associated genes, chromatin profiles, and chromatin states.** A. Density plot showing the global distance of peaks from the nearest transcription start site (TSS), with orange lines indicating a ±1.5 kb distance. B. Distribution of all ATAC-seq peaks among annotated genomic features. C. Number of peaks and associated genes in each aCluster. D. Heatmap of Gene Ontology (GO) terms (Biological Process, BP) for each cluster, with yellow representing more significant enrichment (−log_10_[q-value]). E. Heatmap displaying the similarity (Pearson Correlation Coefficient, PCC) of chromatin modifications between each aCluster. Significance levels are marked as *p < 0.05, **p < 0.01, ***p < 0.001 and ****p ≤ 0.0001. H. Heatmap showing the enrichment of chromatin states within each aCluster. The red box indicates active chromatin states, while the blue box indicates inactive chromatin states. Colors represent the proportion of peaks in each aCluster associated with the indicated chromatin states, with *p < 0.05 determined by Fisher’s exact test. Chromatin states were calculated as previously described^19^.

**Supporting Figure S3 Characterization of enriched proximal CREs and co-occurring CREs across each aCluster and genotype.** A. Bubble plot showing the top 10 ranked TFBMs in each cluster for each genotype. Circle size represents differentially binding scores from TOBIAS^20^, while color represents the log_2_ fold-change between HS and control conditions. B. Radar charts showing all enriched TFBMs in each cluster for each genotype. Circle size represents differential binding scores from TOBIAS, while color indicates the log_2_ fold-change between HS and control conditions. C. Networks illustrating the top 10% co-occurrence pairs of TF families for each genotype calculated by TF-COMB workflow^52^. Node size represents the proportion of each TF family in the network, and edge width indicates the likelihood of co-occurrence. Bold fonts highlight the six TF families with the highest proportions. Red font denotes the TF family that co-occurred exclusively with other TFs in Mp*hsfs* mutants.

**Supporting Figure S4 Characterization of enriched distal CREs and co-occurring CREs across each aCluster and genotype.** A. Bubble plot showing the top 10 ranked TFBMs in each cluster for each genotype. Circle size represents differentially binding scores from TOBIAS, while color represents the log_2_ fold-change between HS and control conditions. B. Radar charts showing all enriched TFBMs in each cluster for each genotype. Circle size represents differential binding scores from TOBIAS, while color indicates the log_2_ fold-change between HS and control conditions. C. Networks illustrating the top 10% co-occurrence pairs of TF families for each genotype. Node size represents the proportion of each TF family in the network, and edge width indicates the likelihood of co-occurrence. Bold fonts highlight the two TF families with the highest proportions.

**Supporting Figure S5 Reanalysis of HSFA1 DAP-seq data across different species.** A. Summary of the number of peaks and genes re-analyzed from Arabidopsis and tomato HSFA DAP-seq data^7,21^. B. Heatmap showing the similarity of CREs across each DAP-seq dataset, measured by the PCC. C. Representative logos of the top five HSFA1-binding motifs with the lowest e-values in Marchantia, tomato, and Arabidopsis. D. Heatmap showing the similarity of HSFA-targeted orthologs across species, measured by the Jaccard index (where 1 indicates identical and 0 indicates completely different). E. Bubble plot showing the enriched (FDR < 0.05) GO terms of orthologs in each species. Note that no enriched terms were found in the AtHSFA4 dataset, so it was excluded from the graph. F.-G. Reanalysis of HS-induced ATAC-seq data in tomato. F. Bubble plot showing the top 10 ranked TFBMs in each cluster for each time point. Circle size represents differentially binding scores from TOBIAS, while color represents the log_2_ fold-change between HS and control conditions. G. Distribution of top-ranked CREs at each time point. Site frequency represents z-scores for the ratio of CRE sites between positive and negative groups, calculated within a range from 1 kb upstream to 0.5 kb downstream of TSSs using a sliding window of 100 bp and a step size of 25 bp. The data in (F.) & (G.) was reanalyzed from a previous publication^7^.

**Supporting Figure S6 Reanalysis of HSFA1 ChIP-seq data across different species, and their function.** A. Peak density plot (±2.0 kb) near TSSs for TuHSFA1 DAP-seq datasets of genes that were up-regulated, down-regulated, or non-responsive under HS. B. Peak density plot (±2.0 kb) near TSSs for MpHSFA1, and AtHSFA1 DAP-seq datasets of genes that were up-regulated, down-regulated, or non-responsive under HS in Mp*hsfa1* or *Athsfa1* (*Atqk*) mutants. The DEGs in Arabidopsis were re-analyzed from previous publications^6^. C. Meta-plots showing HSF1 binding near TSSs between control and HS conditions for each species. D. Peak density plot (±2.0 kb) near TSSs for AtHSFA1, and HsHSFA1 ChIP-seq datasets of genes that were up-regulated, down-regulated, or non-responsive under control or HS conditions. The HS-responsive DEGs in Arabidopsis and human were re-analyzed from previous publications^6,62,63^. E. Venn diagrams show overlapping GO terms identified from HS-upregulated, HSFA1-bound, and HSFA1-unbound orthologous genes across species. F. The heatmap displays the top five enriched GO terms shared among the four species.

**Supporting Figure S7 Characterization of HS-related mutants selected from subnetwork.** A. Bar graph displaying the percentage (%) change in TF expression in Mp*hsfs* mutants compared to Tak-1 plants within the network. ***p < 0.001, determined by hypergeometric distribution. B. Scatter plots showing the expression of all genes within the network between Tak-1 plants and Mp*hsfa1* mutants. *r* indicates the *PCC*. The black and blue lines show the fitted linear regression. C. Subnetworks of filtered nodes. These were selected based on their degree of centrality (bottom 25%) and the number of edges (greater than 25%) in the network. D. Indel information of Mpabi5b-1 and Mpabi5b-12 plants. The red fonts indicate the deletions, and the yellow boxes highlight the gRNAs. B. Representative photos illustrating the morphology of Mp*abi5b-1* and Mp*abi5b-12* plants, compared to Tak-1 plants under control conditions. E. Representative photos of plants after 14 days and 21 days grown on the ½ × B5 plates. F. Thallus area of Mp*abi5b* mutants and Tak-1 plants over time on 1/2 × B5 plates. Dots represent individual plant data, with ***p < 0.001, determined by the one-way ANOVA with Tukey HSD. *n* = 7 plants per line. Boxplots display all data points (dots), median values, first and third quartiles, and whiskers indicating maximum and minimum values. G. Thermotolerance in 7-day-old Tak-1 plants and Mp*wrky3* mutants after a 120-minute HS treatment at 37°C in a water bath. Representative images of plants after five days of recovery are shown in GRB photos. H. Box plots showing the survival rate (%) for each genotype after HS treatment, including the median values, first and third quartiles, and whiskers representing the maximum and minimum values. The p-value was calculated using a two-tailed Student’s t-test; n = 20 gemmalings per genotype across eight independent experiments. The box plots include all data points (dots), median values, first and third quartiles, and whiskers representing the maximum and minimum values. I. PCA analysis of HS-responsive genes in *wrky10*, *abi5b*, and Tak-1 plants.

**Supporting Figure S8 Analysis of RNA-seq and ATAC-seq data in Mp*hsfa1* mutants and Tak-1 plants under ABA treatment.** A & B. Heatmap showing the enriched (FDR < 0.05) GO terms of DEGs (A.) or OCRs (B.) in Mp*hsfa1* mutants under ABA treatment. The numbers in (B.) indicate the count of differential OCRs identified in Mp*hsfa1* mutants.

**Supporting Figure S9 Analysis of HS responses in other species.** A. Heatmap displaying the similarity of HS-responsive differentially expressed orthologs (DEOGs) and differentially expressed orthologous transcription factors (TFOGs) across species, measured by the Jaccard index (where 1 indicates identical and 0 indicates completely different). B. Bubble plot showing the enriched (FDR < 0.05) GO terms of orthologs in each species. C. Heatmaps indicating the presence or absence of each HS-responsive TF family in each species. The red font highlights TF families that also exhibited enriched TFBMs in the CRE analysis.

**Supporting Figure S10 Analysis of ABA responses in other species.** A. Heatmap displaying the similarity of ABA-responsive differentially expressed orthologs (DEOGs) and differentially expressed orthologous transcription factors (TFOGs) across species, measured by the Jaccard index (where 1 indicates identical and 0 indicates completely different). B. Bubble plot showing the enriched (FDR < 0.05) GO terms of orthologs in each species. C. Heatmaps indicating the presence or absence of each HS-responsive TF family in each species.

**Supporting Figure S11 Characterization of enriched CREs in HS- and ABA-responsive genes across different species.** A-D. Bubble plots displaying key features used for model training in HS-responsive (A, C) or ABA-responsive (B, D) genes. Input features are ranked by Cohen’s d effect size (A, C) and importance scores derived from the model (B, D). Higher ranks indicate greater feature importance, with feature colors representing the difference in values between true positives (TPs) and true negatives (TNs), as measured by Cohen’s d effect size. Higher effect size values indicate that the mean feature values of TPs are greater than those of TNs. Circle size reflects the frequency of each CRE across species models. E. Venn diagram showing the overlapping CREs identified in both HS- and ABA-responsive genes.

**Supporting Figure S12 Cross-species and cross-stress prediction using RNA-seq and ATA-seq datasets.** A. Prediction performance of up-regulated and down-regulated genes is represented as AUROC scores in RNA-seq alone or RNA-seq combined with ATAC-seq datasets. Shapes indicate the mean AUROC scores for each ML algorithm, averaged over 10-fold cross-validation. The red dashed line represents the baseline performance expected from random guessing. B. Heatmap showing cross-species and cross-stress prediction performance, using models trained on one species or one stress to predict HS or ABA responses in another, based on RNA-seq alone or RNA-seq combined with ATAC-seq datasets. AUROC scores represent the mean values from 10-fold cross-validation. Full ML results and CRE information are provided in Supporting Dataset S13.

**Supporting Dataset S1 Annotated lists of all ATAC-seq peaks called in different genotypes and conditions at HS**

**Supporting Dataset S2 List of differential open chromatin in aClusters**

**Supporting Dataset S3 List of differentially expressed genes and their enriched cis-regulatory elements in rClusters**

**Supporting Dataset S4 List of enriched cis-regulatory elements in aClusters**

**Supporting Dataset S5 List of co-occurring cis-regulatory elements in aClusters**

**Supporting Dataset S6 List of HSFA1-binding genes from DAP-seq in Marchantia, tomato, *T. uratu,* and Arabidopsis**

**Supporting Dataset S7 List of all genes in the gene regulatory network**

**Supporting Dataset S8 List of differentially expressed genes in *wrky10*, *abi5b*, and Tak-1 plants under HS treatment**

**Supporting Dataset S9 List of differentially expressed genes and open chromatin regions at ABA treatment**

**Supporting Dataset S10 List of expressions and open chromatin regions for 52 ABA bench marker genes**

**Supporting Dataset S11 Raw data for ML results**

**Supporting Dataset S12 List of publicly available re-analyzed datasets**

**Supporting Dataset S13 Raw data for cross-species and cross-stress ML results**

**Supporting Dataset S14 List of primers used in this study**

## Notes

### Competing Interest Statement

The authors have declared no competing interest.

## References

1 Schmitz, R. J. et al. Cis-regulatory sequences in plants: Their importance, discovery, and future challenges. The Plant Cell 34, 718–741 (2021).

2 Misteli, T. & Finn, E. H. Chromatin architecture is a flexible foundation for gene expression. Nature Genetics 53, 426–427 (2021).

3 Wu, T. Y. et al. Diversification of heat shock transcription factors expanded thermal stress responses during early plant evolution. Plant Cell 34, 3557–3576 (2022).

4 Scharf, K. D. et al. The plant heat stress transcription factor (Hsf) family: structure, function and evolution. Biochim Biophys Acta 1819, 104–119 (2012).

5 Fragkostefanakis, S. et al. The repressor and co-activator HsfB1 regulates the major heat stress transcription factors in tomato. Plant Cell Environ 42, 874–890 (2019).

6 Liu, H. C. et al. The role of class A1 heat shock factors (HSFA1s) in response to heat and other stresses in Arabidopsis. Plant Cell Environ 34, 738–751 (2011).

7 Huang, Y. et al. HSFA1a modulates plant heat stress responses and alters the 3D chromatin organization of enhancer-promoter interactions. Nat Commun 14, 469 (2023).

8 Marand, A. P. et al. cis-Regulatory Elements in Plant Development, Adaptation, and Evolution. Annu Rev Plant Biol 74, 111–137 (2023).

9 Gonsalves, S. E. et al. Whole-genome analysis reveals that active heat shock factor binding sites are mostly associated with non-heat shock genes in Drosophila melanogaster. PLoS One 6, e15934 (2011).

10 Mahat, Dig B. et al. Mammalian Heat Shock Response and Mechanisms Underlying Its Genome-wide Transcriptional Regulation. Mol Cell 62, 63–78 (2016).

11 Cortijo, S. et al. Transcriptional Regulation of the Ambient Temperature Response by H2A.Z Nucleosomes and HSF1 Transcription Factors in Arabidopsis. Mol Plant 10, 1258–1273 (2017).

12 Li, B. J. et al. Transcriptional Profiling Reveals a Time-of-Day-Specific Role of REVEILLE 4/8 in Regulating the First Wave of Heat Shock-Induced Gene Expression in Arabidopsis. Plant Cell 31, 2353–2369 (2019).

13 Li, N. et al. Plant Hormone-Mediated Regulation of Heat Tolerance in Response to Global Climate Change. Front Plant Sci 11, 627969 (2020).

14 Ohama, N. et al. Transcriptional Regulatory Network of Plant Heat Stress Response. Trends Plant Sci 22, 53–65 (2017).

15 Bohn, L. et al. The temperature sensor TWA1 is required for thermotolerance in Arabidopsis. Nature 629, 1126–1132 (2024).

16 Albertos, P. et al. Transcription factor BES1 interacts with HSFA1 to promote heat stress resistance of plants. Embo J 41 (2022).

17 Voss, T. C. & Hager, G. L. Dynamic regulation of transcriptional states by chromatin and transcription factors. Nat Rev Genet 15, 69–81 (2014).

18 Montgomery, S. A. et al. Chromatin Organization in Early Land Plants Reveals an Ancestral Association between H3K27me3, Transposons, and Constitutive Heterochromatin. Curr Biol 30, 573-+ (2020).

19 Sequeira-Mendes, J. et al. The Functional Topography of the Arabidopsis Genome Is Organized in a Reduced Number of Linear Motifs of Chromatin States. Plant Cell 26, 2351–2366 (2014).

20 Bentsen, M. et al. ATAC-seq footprinting unravels kinetics of transcription factor binding during zygotic genome activation. Nat Commun 11, 4267 (2020).

21 O’Malley, R. C. et al. Cistrome and Epicistrome Features Shape the Regulatory DNA Landscape. Cell 165, 1280–1292 (2016).

22 Dastidar, S. G. et al. Transcriptional responses of cancer cells to heat shock-inducing stimuli involve amplification of robust HSF1 binding. Nat Commun 14, 7420 (2023).

23 Jahan, A. et al. Archetypal Roles of an Abscisic Acid Receptor in Drought and Sugar Responses in Liverworts. Plant Physiol 179, 317–328 (2019).

24 Sanchez-Olvera, M. et al. ABA-receptor agonist iSB09 decreases soil water consumption and increases tomato CO2 assimilation and water use efficiency under drought stress. Environmental and Experimental Botany 225, 105847 (2024).

25 Jung, H. et al. Nuclear OsFKBP20-1b maintains SR34 stability and promotes the splicing of retained introns upon ABA exposure in rice. New Phytol 238, 2476–2494 (2023).

26 Vendramin, S. et al. Epigenetic Regulation of ABA-Induced Transcriptional Responses in Maize. G3 (Bethesda) 10, 1727–1743 (2020).

27 Seller, C. A. & Schroeder, J. I. Distinct guard cell-specific remodeling of chromatin accessibility during abscisic acid- and CO(2)-dependent stomatal regulation. Proc Natl Acad Sci U S A 120, e2310670120 (2023).

28 Rieseberg, T. P. et al. Time-resolved oxidative signal convergence across the algae–embryophyte divide. bioRxiv, 2024.2003.2011.584470 (2024).

29 Zhang, Y. et al. Evolutionary rewiring of the wheat transcriptional regulatory network by lineage-specific transposable elements. Genome Res 31, 2276–2289 (2021).

30 Lai, W. K. M. & Pugh, B. F. Understanding nucleosome dynamics and their links to gene expression and DNA replication. Nat Rev Mol Cell Biol 18, 548–562 (2017).

31 Guertin, M. J. & Lis, J. T. Chromatin landscape dictates HSF binding to target DNA elements. PLoS Genet 6, e1001114 (2010).

32 Miozzo, F. et al. HSFs, Stress Sensors and Sculptors of Transcription Compartments and Epigenetic Landscapes. Journal of Molecular Biology 427, 3793–3816 (2015).

33 Vihervaara, A. et al. Transcriptional response to stress is pre-wired by promoter and enhancer architecture. Nat Commun 8, 255 (2017).

34 Rao, S. et al. Cooperative binding between distant transcription factors is a hallmark of active enhancers. Mol Cell 81, 1651–1665.e1654 (2021).

35 Kribelbauer, J. F. et al. Low-Affinity Binding Sites and the Transcription Factor Specificity Paradox in Eukaryotes. Annu Rev Cell Dev Biol 35, 357–379 (2019).

36 Balsalobre, A. & Drouin, J. Pioneer factors as master regulators of the epigenome and cell fate. Nat Rev Mol Cell Biol 23, 449–464 (2022).

37 Krishnamoorthi, S. et al. Hyperspectral imaging of liverwort Marchantia polymorpha identifies MpWRKY10 as a key regulator defining Foliar pigmentation patterns. Cell Rep 43, 114463 (2024).

38 Wu, T. Y. et al. G protein controls stress readiness by modulating transcriptional and metabolic homeostasis in Arabidopsis thaliana and Marchantia polymorpha. Mol Plant 15, 1889–1907 (2022).

39 Eklund, D. M. et al. An Evolutionarily Conserved Abscisic Acid Signaling Pathway Regulates Dormancy in the Liverwort Marchantia polymorpha. Curr Biol 28, 3691–3699.e3693 (2018).

40 Gosai, S. J. et al. Machine-guided design of cell-type-targeting cis-regulatory elements. Nature 634, 1211–1220 (2024).

41 Brooks, M. D. et al. Network Walking charts transcriptional dynamics of nitrogen signaling by integrating validated and predicted genome-wide interactions. Nat Commun 10, 1569 (2019).

42 Dündar, G. et al. The heat shock response of plants: new insights into modes of perception and signaling and how hormones contribute. J Exp Bot (2024).

43 Wu, T. Y. et al. Evolutionarily conserved hierarchical gene regulatory networks for plant salt stress response. Nat Plants 7, 787–799 (2021).

44 Kubota, A. et al. Efficient Agrobacterium-mediated transformation of the liverwort Marchantia polymorpha using regenerating thalli. Biosci Biotechnol Biochem 77, 167–172 (2013).

45 Lin, Y. P. et al. Chlorophyll dephytylase 1 and chlorophyll synthase: a chlorophyll salvage pathway for the turnover of photosystems I and II. Plant J 111, 979–994 (2022).

46 Love, M. I. et al. Moderated estimation of fold change and dispersion for RNA-seq data with DESeq2. Genome Biol 15, 550 (2014).

47 Gu, Z. et al. Complex heatmaps reveal patterns and correlations in multidimensional genomic data. Bioinformatics 32, 2847–2849 (2016).

48 Zhang, Y. et al. Model-based analysis of ChIP-Seq (MACS). Genome Biol 9, R137 (2008).

49 Lawrence, M. et al. Software for computing and annotating genomic ranges. PLoS Comput Biol 9, e1003118 (2013).

50 Yu, G. et al. ChIPseeker: an R/Bioconductor package for ChIP peak annotation, comparison and visualization. Bioinformatics 31, 2382–2383 (2015).

51 Ramírez, F. et al. deepTools2: a next generation web server for deep-sequencing data analysis. Nucleic Acids Res 44, W160–W165 (2016).

52 Bentsen, M. et al. TF-COMB – Discovering grammar of transcription factor binding sites. Computational and Structural Biotechnology Journal 20, 4040–4051 (2022).

53 Langmead, B. & Salzberg, S. L. Fast gapped-read alignment with Bowtie 2. Nat Methods 9, 357–359 (2012).

54 Sikora-Wohlfeld, W. et al. Assessing computational methods for transcription factor target gene identification based on ChIP-seq data. PLoS Comput Biol 9, e1003342 (2013).

55 Liu, M.-J. et al. Regulatory Divergence in Wound-Responsive Gene Expression between Domesticated and Wild Tomato. The Plant Cell 30, 1445–1460 (2018).

56 Gupta, S. et al. Quantifying similarity between motifs. Genome Biol 8, R24 (2007).

57 Lachmann, A. et al. ARACNe-AP: gene network reverse engineering through adaptive partitioning inference of mutual information. Bioinformatics 32, 2233–2235 (2016).

58 Huynh-Thu, V. A. et al. Inferring regulatory networks from expression data using tree-based methods. PLoS One 5 (2010).

59 Bailey, T. L. et al. The MEME Suite. Nucleic Acids Res 43, W39–49 (2015).

60 Wu, T. Y. et al. Modeling alternative translation initiation sites in plants reveals evolutionarily conserved cis-regulatory codes in eukaryotes. Genome Res (2024).

61 Emms, D. M. & Kelly, S. OrthoFinder: phylogenetic orthology inference for comparative genomics. Genome Biol 20, 238 (2019).

62 Liu, H., et al. Inflammatory stress-mediated chromatin changes underlie dysfunction in endothelial cells. bioRxiv (2023).

63 Liu, H.-c. & Charng, Y.-y. Common and Distinct Functions of Arabidopsis Class A1 and A2 Heat Shock Factors in Diverse Abiotic Stress Responses and Development Plant Physiology 163, 276–290 (2013).

