## Supporting_figures for "Evolutionarily Conserved Heat-Induced Chromatin Dynamics Drive Heat Stress Responses in Plants"

### Supporting Figure 1

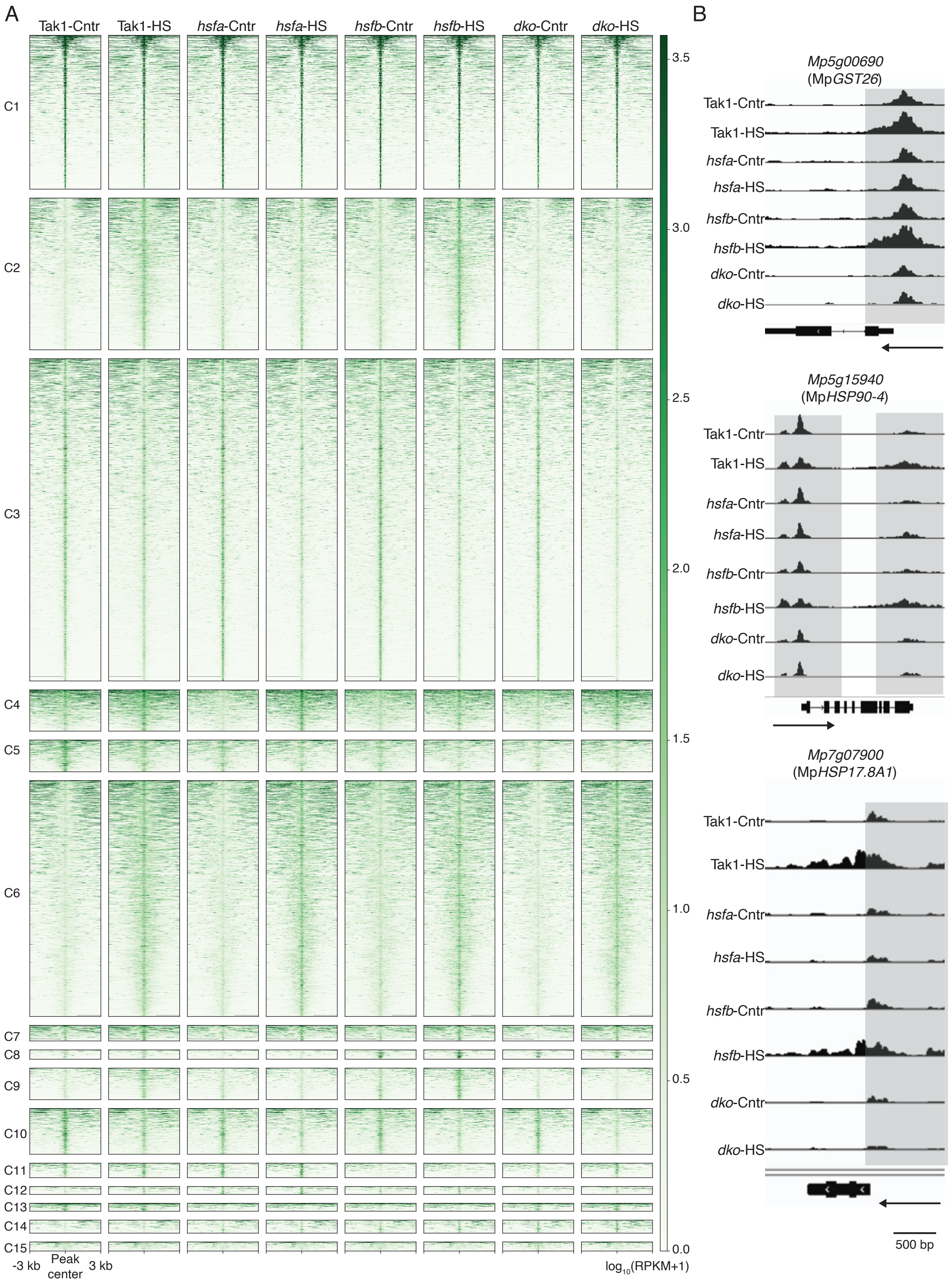

### Supporting Figure 2

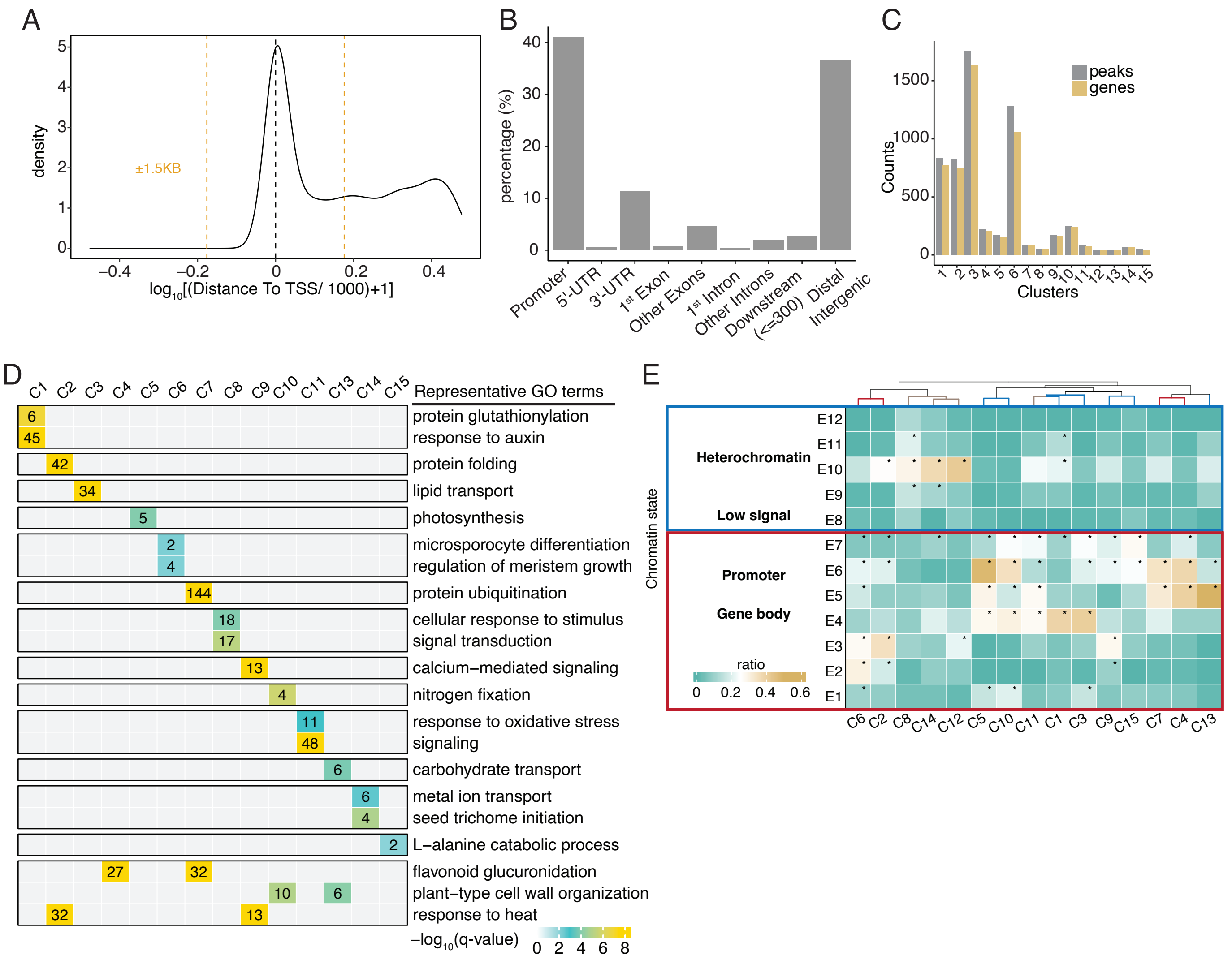

### Supporting Figure 3

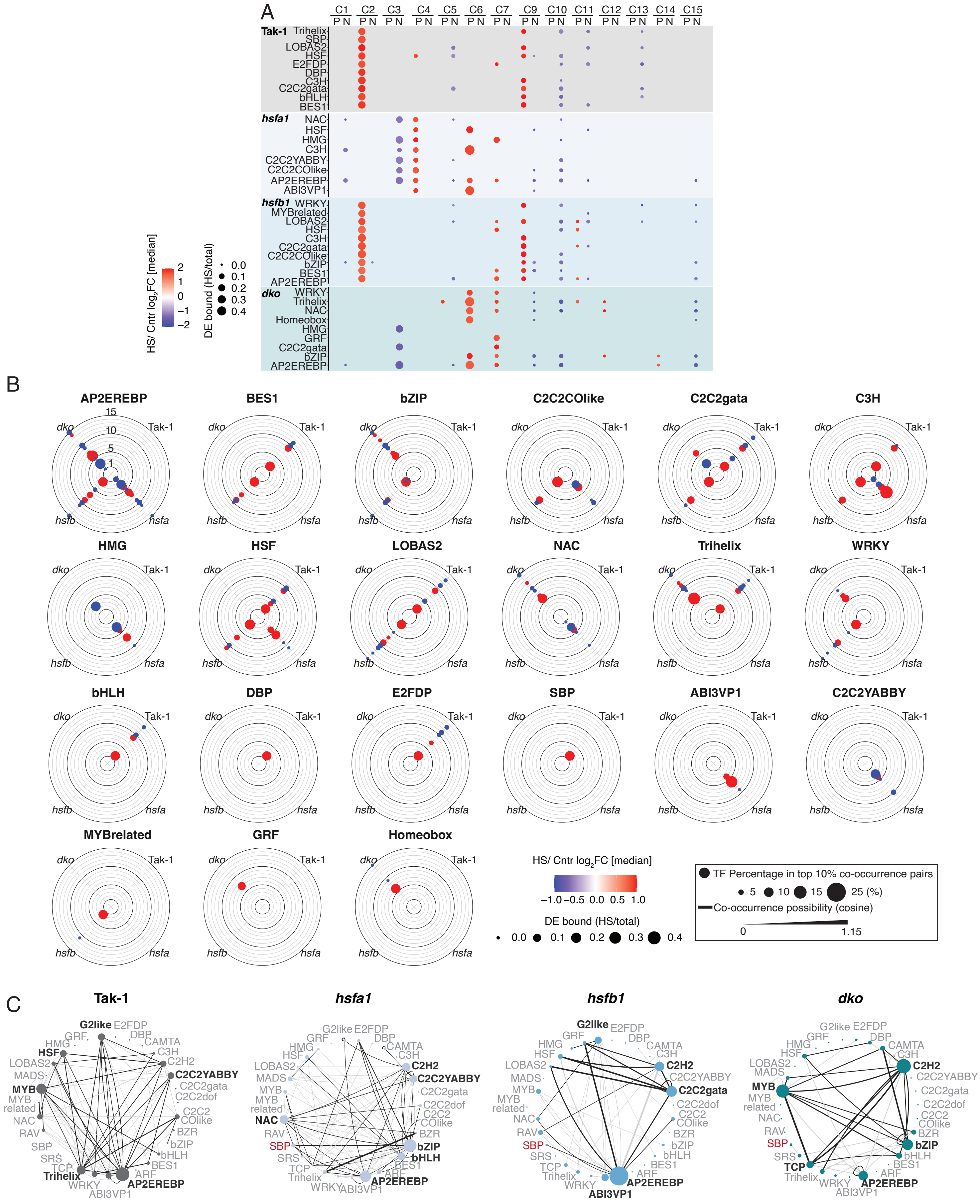

### Supporting Figure 4

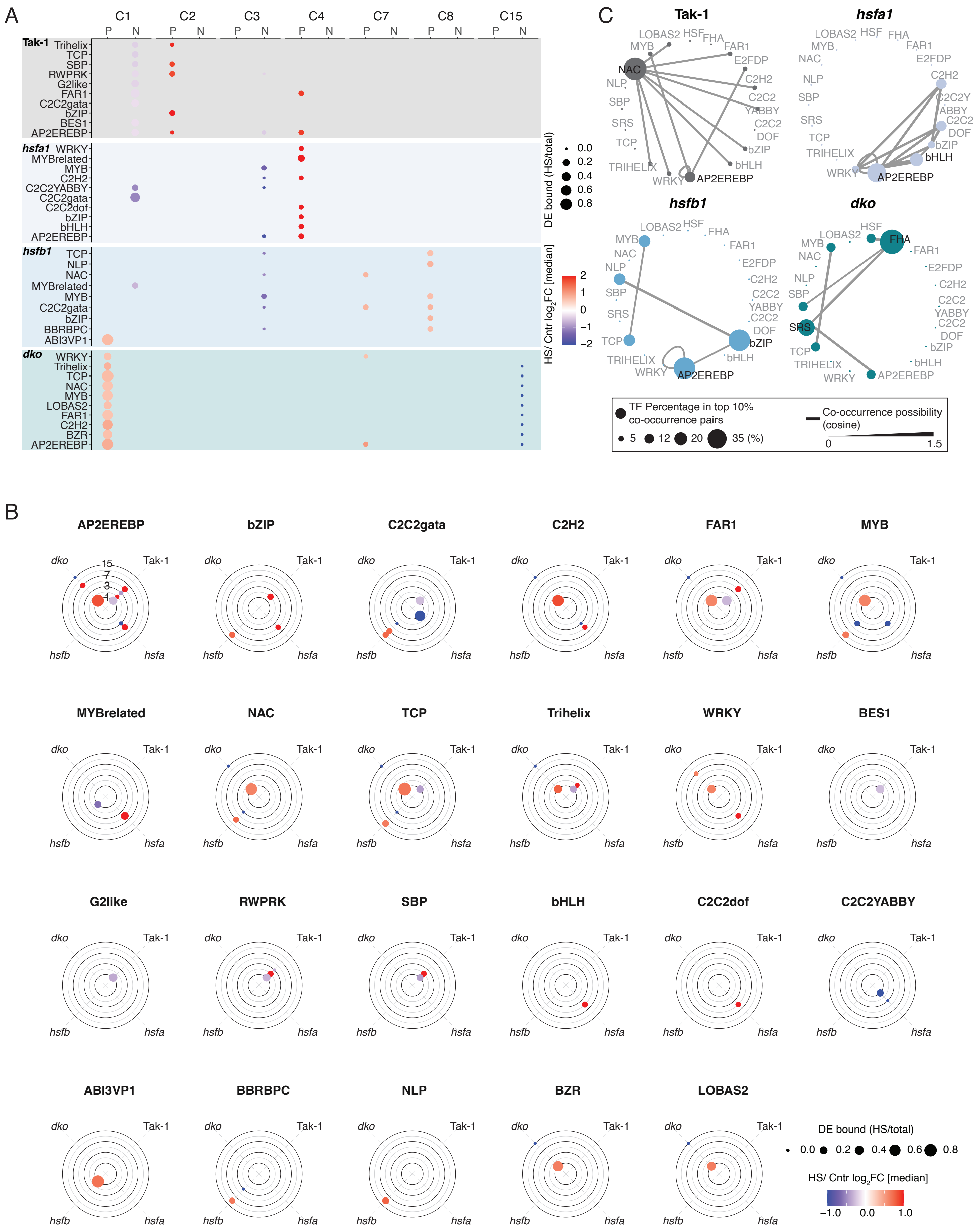

### Supporting Figure 5

A

| ID | Peak | Gene |
|---|---|---|
| MpHSFA1 | 45029 | 5639 |
| SlHSFA1 | 246659 | 6664 |
| AtHSFA1B | 24549 | 7378 |
| AtHSFA1E | 916 | 189 |
| AtHSFA4 | 862 | 291 |
| AtHSFA6A | 1068 | 307 |
| AtHSFA6B | 18478 | 5568 |
| TuHSFA1a | 181920 | 2785 |

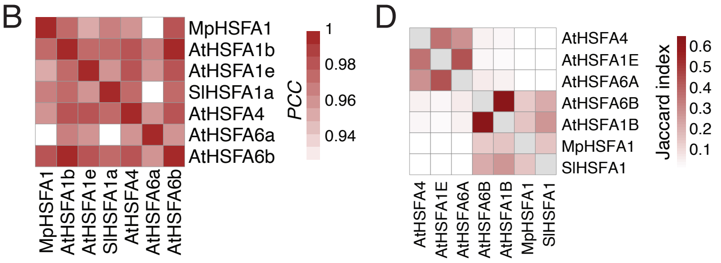


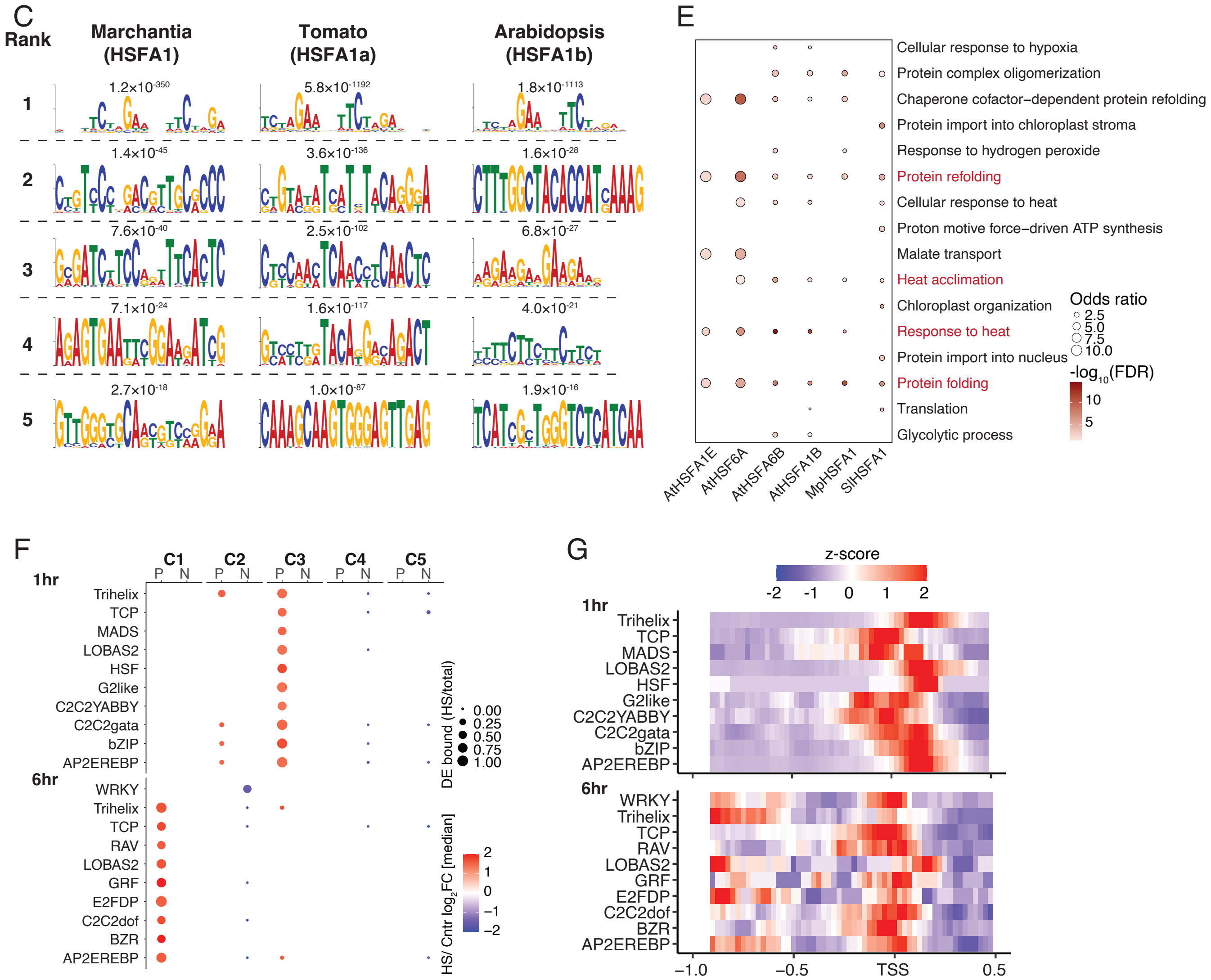

### Supporting Figure 6

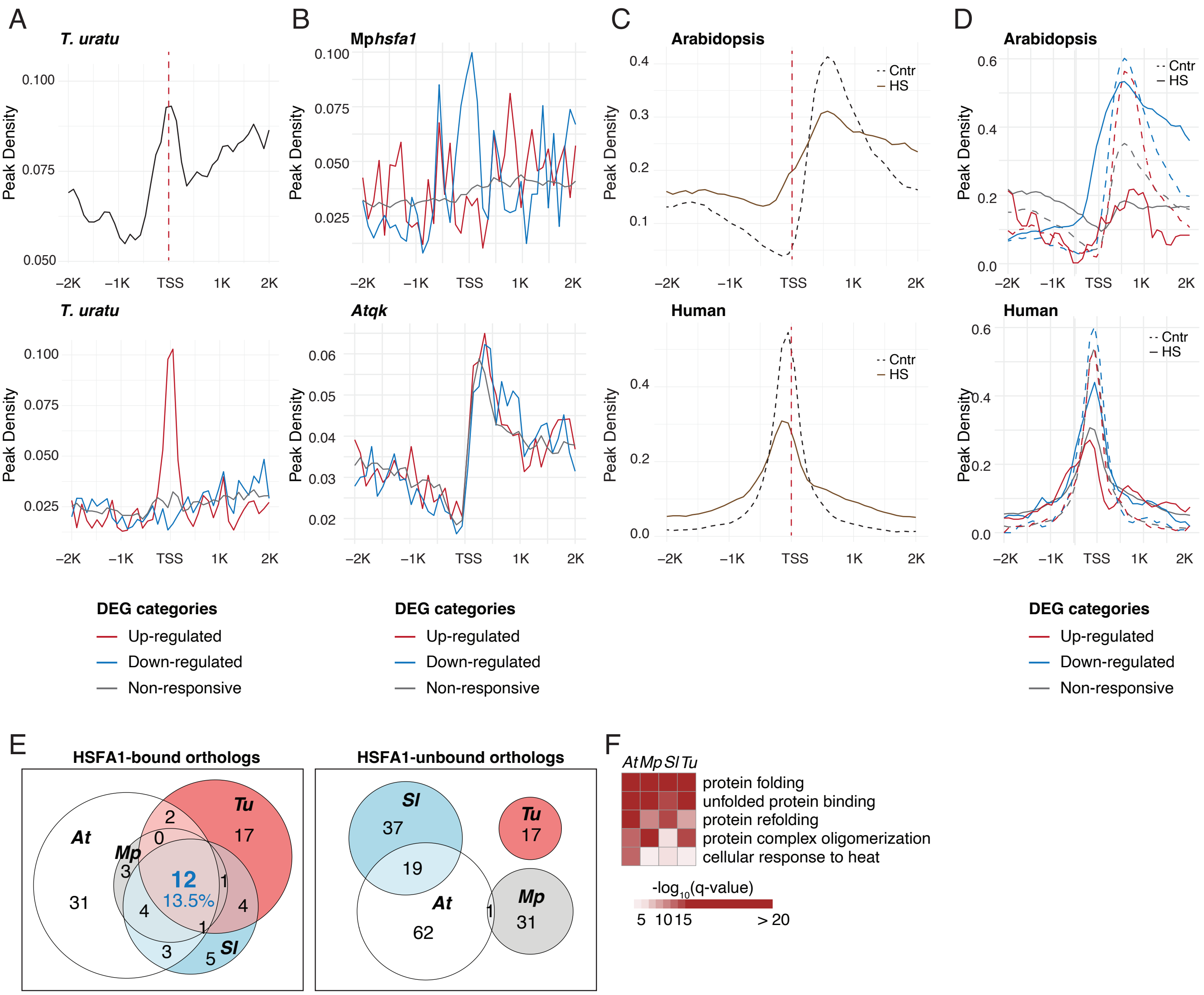

### Supporting Figure 7

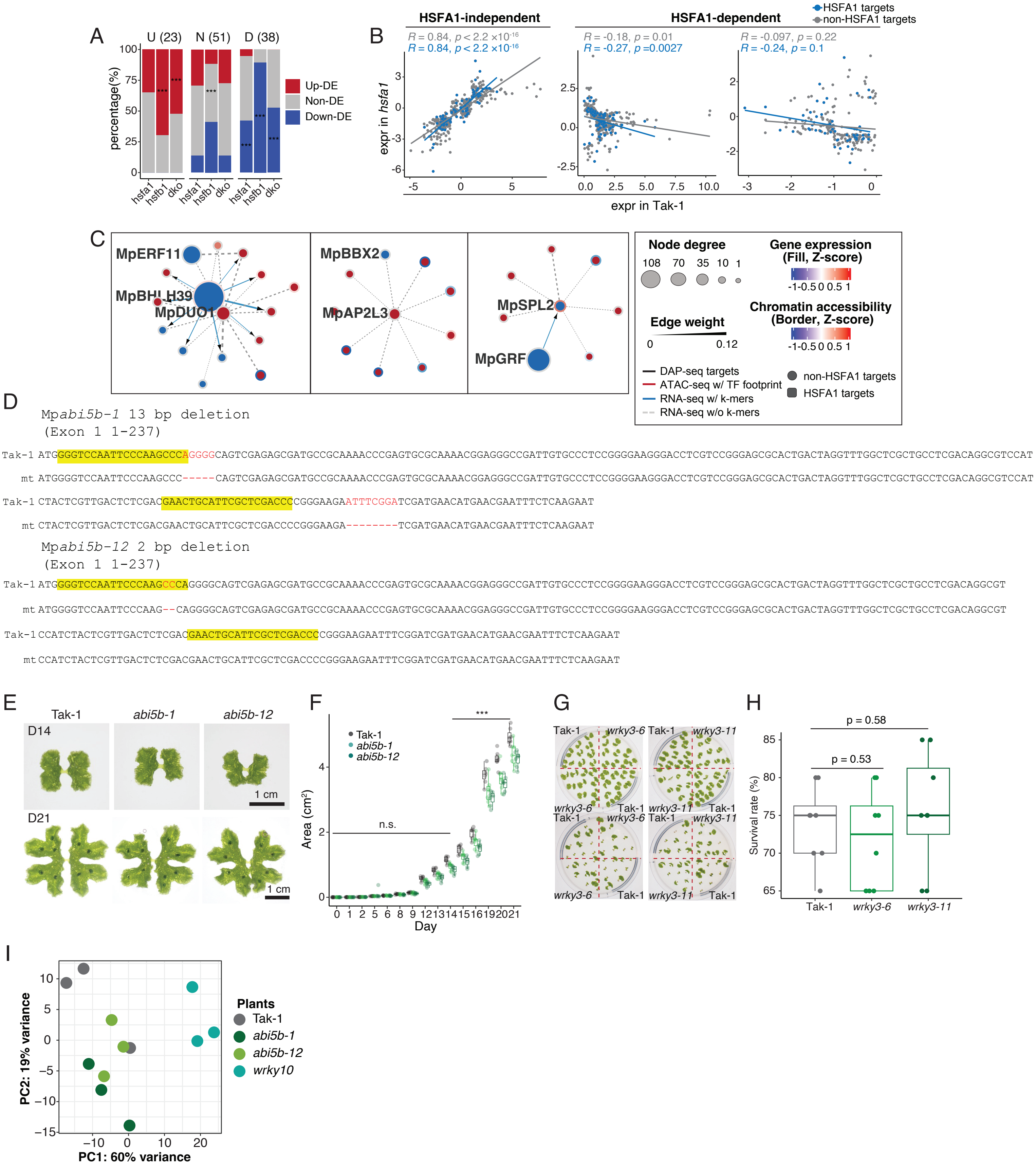

### Supporting Figure 8

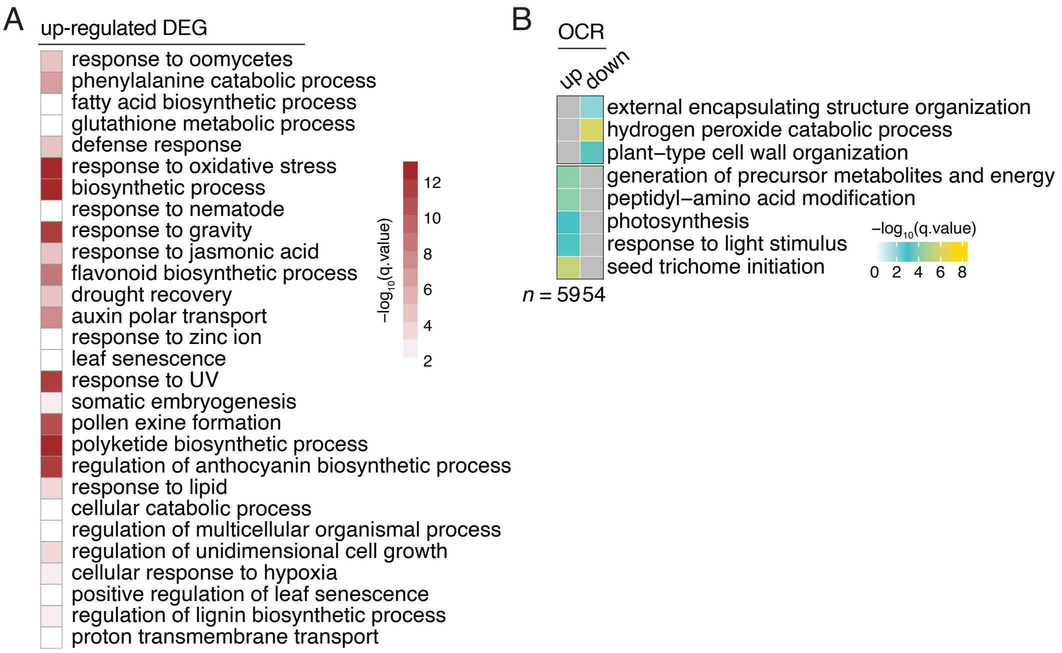

### Supporting Figure 9

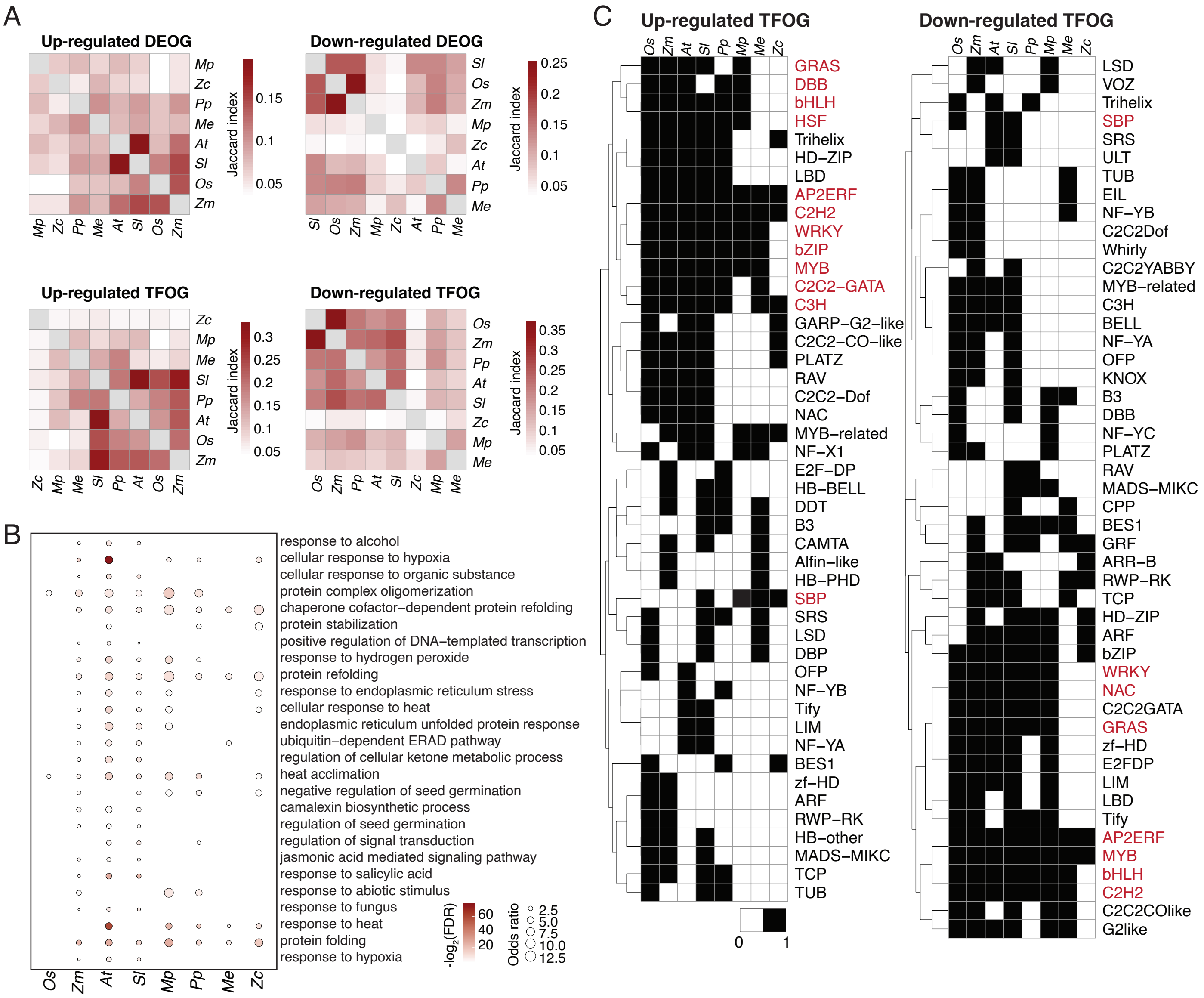

### Supporting Figure 10

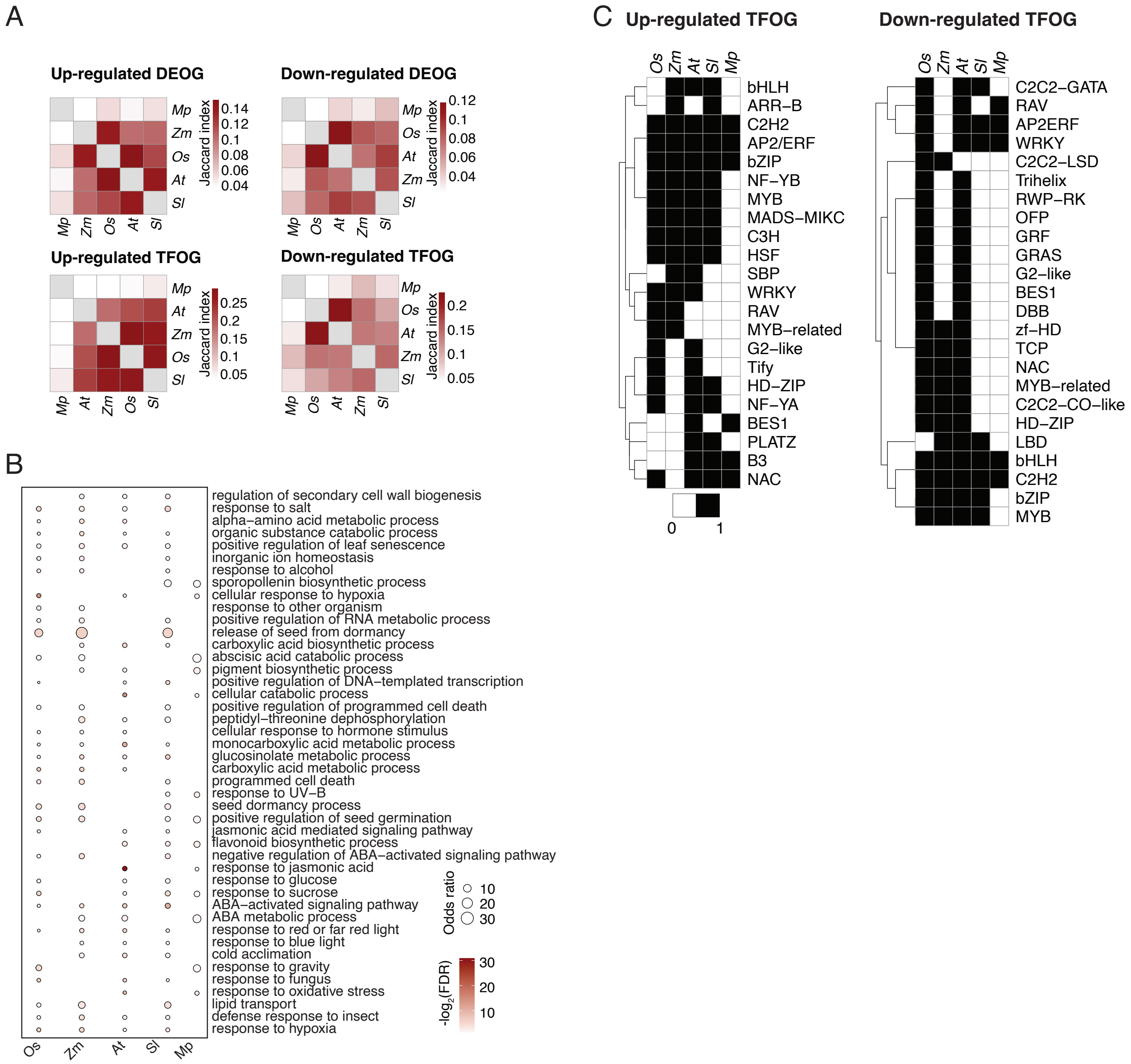

### Supporting Figure 11

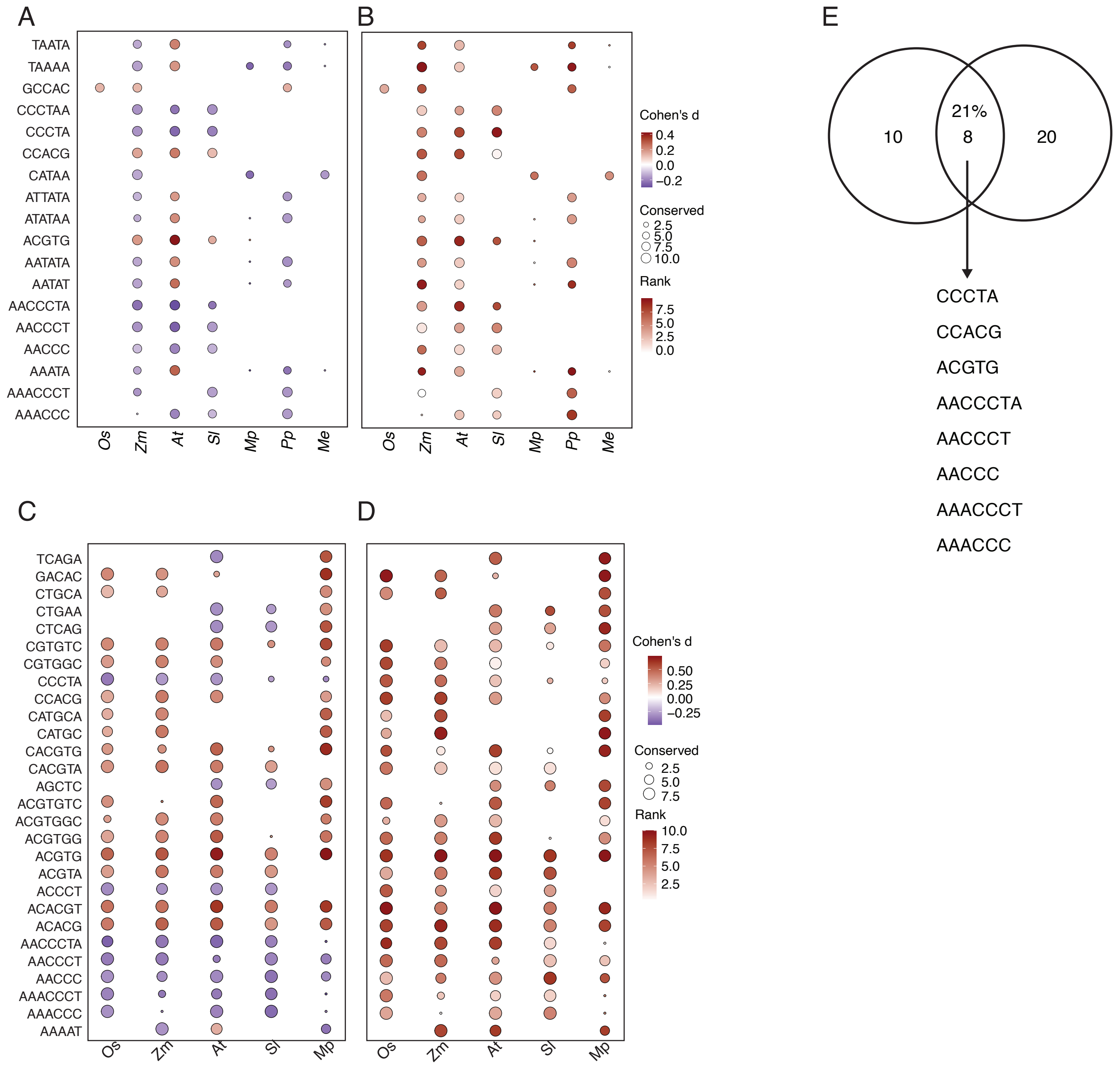

### Supporting Figure 12

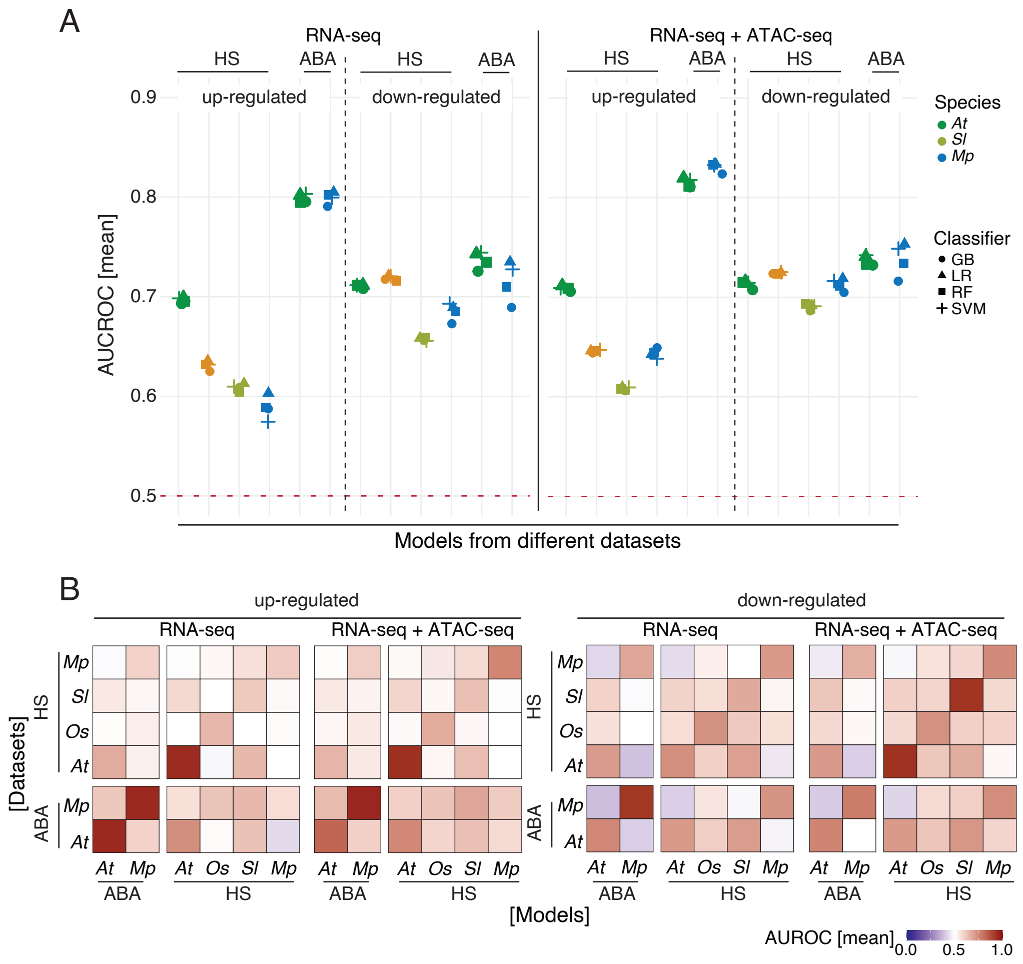
